# At-RS31 orchestrates hierarchical cross-regulation of splicing factors and integrates alternative splicing with TOR-ABA pathways

**DOI:** 10.1101/2024.12.04.626797

**Authors:** Tino Köster, Peter Venhuizen, Martin Lewinski, Ezequiel Petrillo, Yamile Marquez, Armin Fuchs, Debashish Ray, Barbara A. Nimeth, Stefan Riegler, Sophie Franzmeier, Hong Zheng, Timothy Hughes, Quaid Morris, Andrea Barta, Dorothee Staiger, Maria Kalyna

## Abstract

- Alternative splicing is essential for plants, enabling a single gene to produce multiple transcript variants to boost functional diversity and fine-tune responses to environmental and developmental cues. At-RS31, a plant-specific splicing factor in the Serine/Arginine (SR)-rich protein family, responds to light and the Target of Rapamycin (TOR) signaling pathway, yet its downstream targets and regulatory impact remain unknown.
- To identify At-RS31 targets, we applied individual-nucleotide resolution crosslinking and immunoprecipitation (iCLIP) and RNAcompete assays. Transcriptomic analyses of At-RS31 mutant and overexpressing plants further revealed its effects on alternative splicing.
- iCLIP identified 4,034 At-RS31 binding sites across 1,421 genes, enriched in CU-rich and CAGA RNA motifs. Comparative iCLIP and RNAcompete data indicate that the RS domain of At-RS31 may influence its binding specificity *in planta*, underscoring the value of combining *in vivo* and *in vitro* approaches. Transcriptomic analysis showed that At-RS31 modulates diverse splicing events, particularly intron retention and exitron splicing, and influences other splicing modulators, acting as a hierarchical regulator.
- By regulating stress-response genes and genes in both TOR and abscisic acid (ABA) signaling pathways, At-RS31 may help integrate these signals, balancing plant growth with environmental adaptability through alternative splicing.

## INTRODUCTION

Gene expression in eukaryotes involves multiple regulatory layers. Following transcription, nascent RNAs undergo processing steps, including capping, splicing, polyadenylation, and chemical modification, to produce mature mRNAs (Yang *et al*., 2021). Splicing removes introns from pre-mRNA and joins exons to generate the mature transcripts (Gilbert, 1978). Although splicing is a highly regulated process ensuring specificity, it also shows remarkable plasticity. The spliceosome, the cellular machinery responsible for splicing, can recognize alternative splice sites, enabling a single gene to produce multiple transcript variants via alternative splicing.

In plants, 40–70% of intron-containing genes undergo alternative splicing, underscoring its fundamental role in regulating gene expression during development and environmental responses (Filichkin *et al*., 2010; Lu *et al*., 2010; Marquez *et al*., 2012; Chamala *et al*., 2015). Alternative splicing not only produces diverse transcripts leading to different proteins but also generates non-coding isoforms, which may be rapidly degraded or remain stable, thus fine-tuning the total protein levels produced by a gene (Kalyna *et al*., 2012; Petrillo, 2023). Different types of alternative splicing events, such as exon skipping, intron retention, and usage of alternative 5’ and 3’ splice sites, generate transcript diversity. While exon skipping is common in animals, intron retention is most frequent in plants. Retained intron transcripts often remain in the nucleus, regulating protein levels during stress or developmental transitions (Kalyna *et al*., 2012; Marquez *et al*., 2012; Yap *et al*., 2012; Boothby *et al*., 2013; Leviatan *et al*., 2013; Braunschweig *et al*., 2014; Gohring *et al*., 2014; Boutz *et al*., 2015). Furthermore, exitrons, alternatively spliced internal regions within protein-coding exons, add another layer of complexity to the alternative splicing landscape (Marquez *et al*., 2015; Staiger & Simpson, 2015).

The spliceosome ensures the accurate recognition of different pre-mRNA regions and intron removal, aided by numerous proteins. Among these proteins, two key groups stand out: Serine/Arginine-rich (SR) proteins and heterogeneous nuclear ribonucleoproteins (hnRNPs) (Wachter *et al*., 2012). SR proteins interact with the pre-mRNA and spliceosomal components, guiding spliceosome assembly at specific splice sites (Shepard & Hertel, 2009). They contain one or two N-terminal RNA recognition motifs (RRMs), the most prevalent RNA-binding domain, and a C-terminal RS region enriched in arginine/serine dipeptides, which engages primarily in protein-protein interactions but also contributes to RNA recognition. By interacting with spliceosomal machinery, SR proteins modulate splice sites selection, contributing mRNA isoform diversity. Beyond splicing, SR proteins influence transcription (Lin *et al*., 2008; Ji *et al*., 2013), polyadenylation (Schwich *et al*., 2021), mRNA export (Müller-McNicoll *et al*., 2016; Botti *et al*., 2017), and translation (Sanford *et al*., 2004) among other processes. However, most knowledge about SR protein functions comes from animal studies.

In plants, the SR protein family has expanded remarkably, with *Arabidopsis thaliana* possessing 18 SR proteins classified into six subfamilies. Ten are plant-specific, divided into RS, RS2Z, and SCL subfamilies based on their domain organization. The remaining eight are similar to mammalian SR proteins SF2/ASF/SRSF1, 9G8/SRSF7, and SC35/SRSF2 and belong to SR, RSZ, and SC subfamilies, respectively (Kalyna & Barta, 2004; Barta *et al*., 2008; Barta *et al*., 2010; Duque, 2011; Richardson *et al*., 2011). Arabidopsis also has two SR-like proteins, SR45 and SR45a. These proteins participate in constitutive and alternative splicing and play roles in mRNA export, stability, translation, transcriptional elongation, and cell cycle regulation (Jin, 2022). Several SR proteins contribute to plant development and abiotic stress responses (Jin, 2022), but their *in vivo* targets and regulatory networks remain less characterized (Mateos & Staiger, 2022). So far, RNA immunoprecipitation (RIP) followed by RNA sequencing (RNA-seq) identified over 4000 RNAs associated with SR45 in Arabidopsis seedlings (Xing *et al*., 2015) and 1812 in inflorescences (Zhang, X-N *et al*., 2017). Recently, tomato RS2Z35 and RS2Z36, were shown to bind to transcripts of over 5000 genes, including the heat shock transcription factor and many transcripts that undergo heat shock-sensitive alternative splicing, preferentially binding purine-rich RNA motifs (Rosenkranz *et al*., 2024).

At-RS31 (AT3G61860), a plant-specific SR protein in the RS subfamily, may regulate unique plant functions, though its exact roles are unclear. It has two N-terminal RRMs (see Fig. **3a**). RRM1 contains the conserved RDAEDA region like human SF2/ASF/SRSF1, while the RRM2 lacks the typical SWQDLKD motif, diverging from non-plant SR proteins. The RS region has DYGRRPSP repeats, specific to this subfamily (Lopato *et al*., 1996). At-RS31 interacts with SR and SR-like proteins (At-RSZ21, At-RS2Z33, SR45a), spliceosome components (U1-70K, U11/U12-31K), and cyclophilin AtCyp59, suggesting a role in pre-mRNA splicing (Lopato *et al*., 2002; Lorkovic *et al*., 2005; Gullerova *et al*., 2006; Altmann *et al*., 2020). Its ability to stimulate splicing in SR protein-deficient HeLa cell S100 extract further supports this role (Lopato *et al*., 1996).

*At-RS31* undergoes alternative splicing, producing three main transcript isoforms: mRNA1-3 (Lopato *et al*., 1996) (see Fig. **3b**). The shortest isoform, mRNA1, arises from excision of the entire intron 2 and encodes the SR protein. mRNA3 uses a proximal 3’ splice site in intron 2, while mRNA2 arises either from the removal of a small intron in mRNA3 or inclusion of a cassette exon compared to mRNA1. Both mRNA2 and mRNA3 contain premature termination codons (PTCs), but only mRNA2 is sensitive to nonsense-mediated mRNA decay (NMD), likely due to nuclear retention of mRNA3 (Kalyna *et al*., 2012; Petrillo *et al*., 2014).

The ratio of *At-RS31* isoforms varies with tissue type, developmental stage, and environmental stimuli like bacterial flagellin, cold, or red light (Lopato *et al*., 1996; Palusa *et al*., 2007; Tognacca *et al*., 2019; Bazin *et al*., 2020). Proportion of mRNA1 fluctuates in response to light, increasing in light and decreasing in darkness, a response mediated by chloroplast retrograde signalling, affecting even non-photosynthetic root cells lacking chloroplasts; sugars, mitochondrial function, and the Target of Rapamycin (TOR) pathway are key to this effect (Petrillo *et al*., 2014; Riegler *et al*., 2021). At-SR30, At-RS2Z33 and SR45 modulate *At-RS31* alternative splicing (Lopato *et al*., 1999; Kalyna *et al*., 2006; Ali *et al*., 2007; Simpson *et al*., 2012; Carvalho *et al*., 2016). The conserved alternative splicing pattern of *At-RS31* across diverse plant species, from green algae to flowering plants, underscores its biological significance (Iida & Go, 2006; Kalyna *et al*., 2006).

Despite the identification of many factors regulating *At-RS31* alternative splicing, hinting at its potential roles in several plant biological processes, its downstream targets remain unknown. Since SR proteins influence alternative splicing in a concentration-dependent manner (Mayeda *et al*., 1992) and given the dynamic modulation of *At-RS31* in response to various environmental and developmental signals, we hypothesize that its expression levels significantly impact the transcriptome. To identify direct targets of At-RS31, we performed individual-nucleotide resolution crosslinking and immunoprecipitation (iCLIP) (Konig *et al*., 2010; Hafner *et al*., 2021) followed by RNAcompete validation (Ray *et al*., 2013; Ray *et al*., 2017). Additionally, we performed transcriptome profiling in *rs31-1* mutant plants and plants constitutively overexpressing At-RS31 (Petrillo *et al*., 2014). By identifying At-RS31 direct targets through iCLIP and transcripts that are differentially alternatively spliced in response to altered At-RS31 levels, we gain insights into the downstream processes controlled by At-RS31.

## MATERIALS AND METHODS

### Generation of GFP-tagged *At-RS31* genomic construct and transgenic plants for iCLIP

To generate *RS31::RS31-GFP* construct for iCLIP (Fig. **S1a**), the *At-RS31* genomic region including its own promoter, untranslated regions and introns was amplified from *A. thaliana* DNA by PCR. The construct was cloned into pGreenII0029. C-terminal fusion to GFP was achieved by mutating the *At-RS31* reference stop codon *via* site-directed PCR mutagenesis. The resulting construct was used to generate transgenic *A. thaliana* plants *via* floral dipping (Clough & Bent, 1998). *RS31::RS31-GFP* construct was introduced into the *rs31-1* mutant line (see Fig. **3b**). Wild type plants were transformed with *35S::GFP* construct, selected on 50 µg/L kanamycin and by genotyping. Transgenic plants containing *RS31::RS31-GFP* were selected *via* fluorescence stereomicroscopy and genotyping. Genomic DNA was isolated from a single leaf as described by (Edwards *et al*., 1991) and genotyping was performed using primers listed in Table **S1**.

Transgenic plants were assessed for expression of the At-RS31-GFP fusion protein. Plant tissues were ground in liquid nitrogen and resuspended in PEB400 buffer (50 mM HEPES-KOH pH 7.9, 400 mM KCl, 2.5 mM MgCl_2_, 1 mM EDTA pH 8.0, 1 mM DTT, 0.1% Tween-20, protease and phosphatase inhibitor). The crude extract was incubated on ice for 15 min before sonication, followed by three rounds of centrifugation. The final supernatant was diluted with PEB KCl-free buffer to adjust the KCl concentration to 200 mM). Western blotting was performed according to standard procedures. Proteins were separated by 10-16% SDS-PAGE. Immunoblotting (Fig. **S1b**) was performed using rabbit monoclonal anti-GFP (D5.1) antibody (Cell Signaling Technology) at 1:1000 dilution.

### Growth of plant material for iCLIP

Seeds of *RS31::RS31-GFP* and *35S::GFP* plants were sown on agar plates containing half strength Murashige and Skoog (MS) medium: 2.2 g/L MS (Duchefa M0222.0050), 0.5 g/L 2-morpholinoethanesulfonic acid (MES), 1% sucrose, and 1% agar. The plates were placed vertically in growth cabinets and incubated under 16 h/8 h light/dark cycle at 22 °C and 60% humidity for 14 days.

### iCLIP

*RS31::RS31-GFP* and *35S::GFP* seedlings were subjected to irradiation with 254 nm UV-light at a dose of 2000 mJ/cm^2^ (UVP CL-1000 UV crosslinker) on ice 4 h after lights on. iCLIP was performed essentially as described (Meyer *et al*., 2017; Köster & Staiger, 2021). Adapter sequences are provided in Table **S1**.

### Bioinformatics evaluation of iCLIP reads

The quality of sequenced reads was examined using fastqc 0.11.5 (http://www.bioinformatics.bbsrc.ac.uk/projects/fastqc). Adapters at the 3’ end were trimmed using cutadapt version 1.16 (Martin, 2011). The samples were then demultiplexed with the flexbar toolkit version 3.4.0 by applying the additional -bk parameter to conserve the barcode information for further steps (Roehr *et al*., 2017). Reads shorter than 24 nt were discarded. Barcodes were manually trimmed and saved to the *read_id* field. The processed reads were mapped to the TAIR10 genome with STAR version 2.6.0a allowing a maximum of two mismatches and soft clipping only at the 3’ end (Dobin *et al*., 2013). The removal of PCR duplicates was done by grouping the reads by their mapping start position. Reads with the identical start position and random barcode were removed from the samples (Python3 and pybedtools) (Dale *et al*., 2011). The peak calling of uniquely mapped reads was done using PureCLIP 1.0.4 (Krakau *et al*., 2017). The PureCLIP parameters were set to the second profile option (-bc 1) allowing broader regions with less read starts to be called. To remove redundancy after the peak calling, peaks located at immediately adjacent nucleotides were grouped together and only the peak with the highest PureCLIP score was kept. The called peak position was extended by 4 nt upstream and downstream each to define a binding site of 9 nt. The center position of the binding site marks the binding site peak. The extension of the binding site peak positions was computed using bedtools version 2.27.1 (Quinlan & Hall, 2010).

### Motif discovery

The sequence at each binding site (length of 9 nt) was extracted by applying the getfasta program from bedtools (Quinlan & Hall, 2010) in a strand-specific mode (-s) using TAIR10 as the reference genome. A *de novo* motif discovery was then applied using STREME 5.3.3 (Bailey, 2021) to identify significantly enriched motifs with a length between 3 and 8 nucleotides in the binding site sequences. Additionally, *k*-mer distributions with the identical set of binding site sequences were computed using k = 6 (hexamers) similar to the length of significant motifs identified by STREME. To account for local sequence bias, the hexamer counts were normalized by sampling sequences randomly 1000 times from the identical transcripts targeted by the RS31-GFP protein. The randomized set was used to calculate a *z*-score for each hexamer by subtracting the mean of random occurrences from the actual occurrence and dividing it by the standard deviation of the random occurrences.

### Binding site distribution in protein-coding genes

Each binding site was assigned to the overlapping transcript and gene feature, namely 5’UTR, CDS, intron, or 3’UTR according to the Araport11 annotation. Only the representative models from TAIR of a given gene were considered to avoid ambiguity in the assignment. The distribution of binding sites among gene features from RS31::RS31-GFP and 35S::GFP samples were compared using the percentage of mapped binding sites.

To determine the location of At-RS31 binding sites relative to the 5′ splice site, the peak positions of all binding sites were compared against 5’ splice sites from protein-coding genes annotated in Araport11 using a custom R-Script and the *tidyverse* package (https://www.tidyverse.org/). Only exons with a length of at least 50 nucleotides were considered. Sequences around At-RS31 binding sites 20 to 35 nt upstream of 5’ splice sites were extracted using getfasta from bedtools (Quinlan & Hall, 2010) and aligned using the ClustalO Web Service from Jalview 2.11.1.7 (Waterhouse *et al*., 2009). Based on the resulting alignments, a sequence logo was created using WebLogo3 (Crooks *et al*., 2004).

The positional relationship of At-RS31 binding sites to transcription start sites (TSS) was mapped though intersecting the annotated genomic TSS locations (Zhang *et al*., 2022) with genomic coordinates of At-RS31 iCLIP targets using bedtools. The most 5’ TSS for each gene was chosen with a custom R script and distribution of At-RS31 binding site peaks up to 250 nucleotides downstream of the TSS locations was plotted with the ggplot2 R package.

### Generation of GST-tagged At-RS31 constructs and protein purification for RNAcompete assay

Cloning and protein purification were performed as described previously (Ray *et al*., 2017). The full-length At-RS31 construct (T7::GST-31FL) and RRMs-only construct (T7::GST-31RRMs), containing the first 175 amino acids, were generated from *A. thaliana* cDNA by PCR. An artificial stop codon was introduced by site-directed PCR mutagenesis. The constructs were cloned into pTH6838 (Ray *et al*., 2013), downstream of the T7 promoter and fused N-terminally to GST (Fig. **S2a**). Primers are listed in Table **S1**.

GST-tagged proteins were expressed in *E.coli* BL21 cells grown at 37 °C, lysed by sonication in PBS with protease inhibitor (Roche), and purified using glutathione sepharose 4B beads (GE Healthcare). Proteins were eluted, and their concentration and size were assessed by spectrophotometry and SDS-PAGE (Fig. **S2b**).

### RNAcompete assay

RNAcompete assay was performed using purified GST-At-RS31 and GST-At-RS31-RRMs fusion proteins for pull down experiments. RNAcompete analysis was performed essentially as described in (Ray *et al*., 2013; Ray *et al*., 2017).

### RNAcompete CAGA motif statistics

To compare the At-RS31 motifs retrieved by RNAcompete with the iCLIP binding sites, the TAIR10 genome was scanned for positions matching the RNAcompete consensus motif CAGA using FIMO version 4.11.1 (Grant *et al*., 2011). The RS31-GFP binding site peaks were compared directly to the positions of CAGA sites with a custom R script.

### Plant material for RNA sequencing

We used three *A. thaliana* lines: the *At-RS31* overexpression line (*35S::RS31*, overexpressing the protein-coding isoform AT3G61860.1 under the CaMV 35S promoter), the *rs31-1* mutant (SALK_021332, T-DNA insertion in exon 5 of *At-RS31*) (Petrillo *et al*., 2014), and the wild type Col-0 (see Fig. **3b-d**). For each genotype, plants were grown in three biological replicates under the same conditions described for the iCLIP experiments.

### RNA isolation and RNA sequencing

Total RNA was isolated using RNeasy Plant Mini Kit and treated with DNAse (both Qiagen). Strand-specific transcriptome libraries were sequenced at 100 bp paired-end using the Illumina Hi-seq 2000 system (Next Generation Sequencing Facility, Vienna BioCenter) yielding 30-40 G bases for each biological replicate.

### Transcript quantification

The transcript per million (TPM) expression was estimated with Salmon with the additional -- gcBias and --validateMappings flags (version 0.14.1; (Patro *et al*., 2017)) for the Reference Transcript Dataset for Arabidopsis 2 (AtRTD2)-Quantification of Alternatively Spliced Isoforms (QUASI) (AtRTD2-QUASI) annotation (Zhang, R *et al*., 2017).

### Read alignment with STAR

Reads were mapped to the index based on the TAIR10 genome release and the AtRTD2 transcriptome with STAR (version 2.7.1a; (Dobin *et al*., 2013)) using a 2-pass mapping. The following parameters were used: --outSAMprimaryFlag AllBestScore, -- outFilterMismatchNmax 2/0 (first/second pass), --outSjfilterCountTotalMin 10 5 5 5, -- outFilterIntronMotifs RemoveNoncanonical, --alignIntronMin 60, --alignIntronMax 6000, -- outSAMtype BAM SortedByCoordinate. During the second pass, the splice junction files of the control and test samples were passed to the mapping via the --sjdbFileChrStartEnd flag.

### Differential gene expression analysis

Differential expression (DE) analysis was performed using 3D RNA-seq App (Guo *et al*., 2020). Read counts and transcript per million reads (TPMs) were generated using tximport R package version 1.10.0 and lengthScaledTPM method (Soneson *et al*., 2016) based on transcript quantifications from Salmon (Patro *et al*., 2017). Lowly expressed transcripts and genes were filtered by analysing mean-variance trend of the data. Transcripts were considered expressed if they had counts per million (CPM) ≥ 1 in at least 3 of the 9 samples, providing an optimal filter for low expression. A gene was considered expressed if any of its transcripts met the above criteria. The TMM method was used to normalise gene and transcript read counts to log2-CPM (Bullard *et al*., 2010). Limma R package was used for 3D expression comparison (Law *et al*., 2014; Ritchie *et al*., 2015). Expression changes were compared between *rs31-1 vs*. wild type and *35S::RS31 vs.* wild type contrast groups. For DE genes, the log2 fold change (L2FC) of gene abundance was calculated, and significance was determined using a t-test. P-values of multiple testing were adjusted for false discovery rate (FDR) (Benjamini & Yekutieli, 2001). Genes were considered significantly differentially expressed if they had an adjusted p-value < 0.05 and |L2FC| ≥ 1.

### Differential alternative splicing analysis

Alternative splicing events were obtained and quantified using Whippet, version 0.11 (Sterne-Weiler *et al*., 2018). Two separate splice graph indices were generated: one for exon skipping (ES), alternative acceptor (AA) and alternative donor (AD) events, and another for retained introns (RI) and exitrons (EI). Both indices were based on the AtRTD2 transcriptome annotation (Zhang, R *et al*., 2017), supplemented with the STAR RNA-seq alignments, and generated with the additional --bam-min-reads 10 flag. The RI/EI index was further supplemented with “pre-mRNA” gene coordinates and exitron splice junctions detected using an in-house script. “Pre-mRNA” coordinates ranged from the start to the end of genes, allowing quantification the retention levels of all annotated introns in a gene. The Whippet delta step was run with default parameters, except for the --min-samples 3 flag. AA and AD events were filtered to ensure both (alternative) junctions were detected in Whippet data. RI events were required to have at least one read in either all control and/or test samples. Events with probability ≥ 0.9 and absolute delta percent-spliced-in (IPSII) ≥ 0.1 were considered significant differential alternative splicing events.

### Functional enrichment analysis

Functional gene set enrichment analysis was performed using g:GOSt tool of the g:Profiler (Kolberg *et al*., 2023) (annotation version e110_eg57_p18_4b54a898) at https://biit.cs.ut.ee/gprofiler/gost.

### Statistical analysis of gene overlaps and Venn diagrams

The intersections of gene lists were calculated and visualized using the Venn diagram tool available at http://bioinformatics.psb.ugent.be/webtools/Venn/. The statistical significance of gene overlaps was assessed using Fisher’s exact test.

### RT-PCR

One microgram of DNA-free total RNA was reverse transcribed using Avian Myeloblastosis Virus (AMV) reverse transcriptase and oligo(dT)_15 primers (Reverse Transcription System Kit, Promega). PCR amplification was performed with Phusion DNA Polymerase (LifeTechnologies) for 30-35 cycles. *UBQ1* was amplified as a loading control using 25 cycles. Primers are listed in Table **S1**.

### Visualization of transcript isoform schematics

Transcript models for the genes are visualized using Boxify tool (Riegler *et al*., 2021) available at https://boxify.boku.ac.at/.

## RESULTS

### Genome-wide determination of At-RS31 target transcripts and its binding motifs by iCLIP

To identify transcripts bound by At-RS31 *in vivo* across the transcriptome, we utilized iCLIP (Meyer *et al*., 2017; Arribas-Hernandez *et al*., 2021). *RS31::RS31-GFP* plants expressing At-RS31 fused to GFP under control of the endogenous promoter in the *rs31-1* background and *35S::GFP* control plants expressing GFP alone under control of the 35S CaMV promoter (Fig. **S1**) were subjected to UV crosslinking to preserve *in vivo* RNA-protein interactions. After cell lysis, the RNA-protein complexes were immunoprecipitated with GFP Trap beads, and the co-precipitated RNAs were radiolabelled. Upon denaturing gel electrophoresis, membrane transfer and autoradiography, RNA-protein complexes were detected in the crosslinked *RS31::RS31-GFP* plants (Fig. **1a**, left). RNase I treatment strongly reduced the signal, indicating that indeed RNA was crosslinked. For plants expressing GFP alone, only little RNA was cross-linked. The identity of the proteins was confirmed by probing the membrane with an anti-GFP antibody (Fig. **1a**, right). The membrane region corresponding to the covalently linked RS31-GFP–RNA complexes was excised (Fig. **S3a**). Similarly, we excised the region above the GFP band (Fig. **S3b**). RNA was isolated for preparation of libraries for high throughput sequencing from three biological replicates (Fig. **S3c, d**). The read statistics are presented in Table **S2**. As reverse transcriptase stops at the remnants of the digested protein, the binding sites can be retrieved with single nucleotide resolution. A global view of the localization of the crosslink sites of RS31-GFP and GFP alone throughout the chromosomes is shown in Fig. **S4**.

**Fig. 1.**
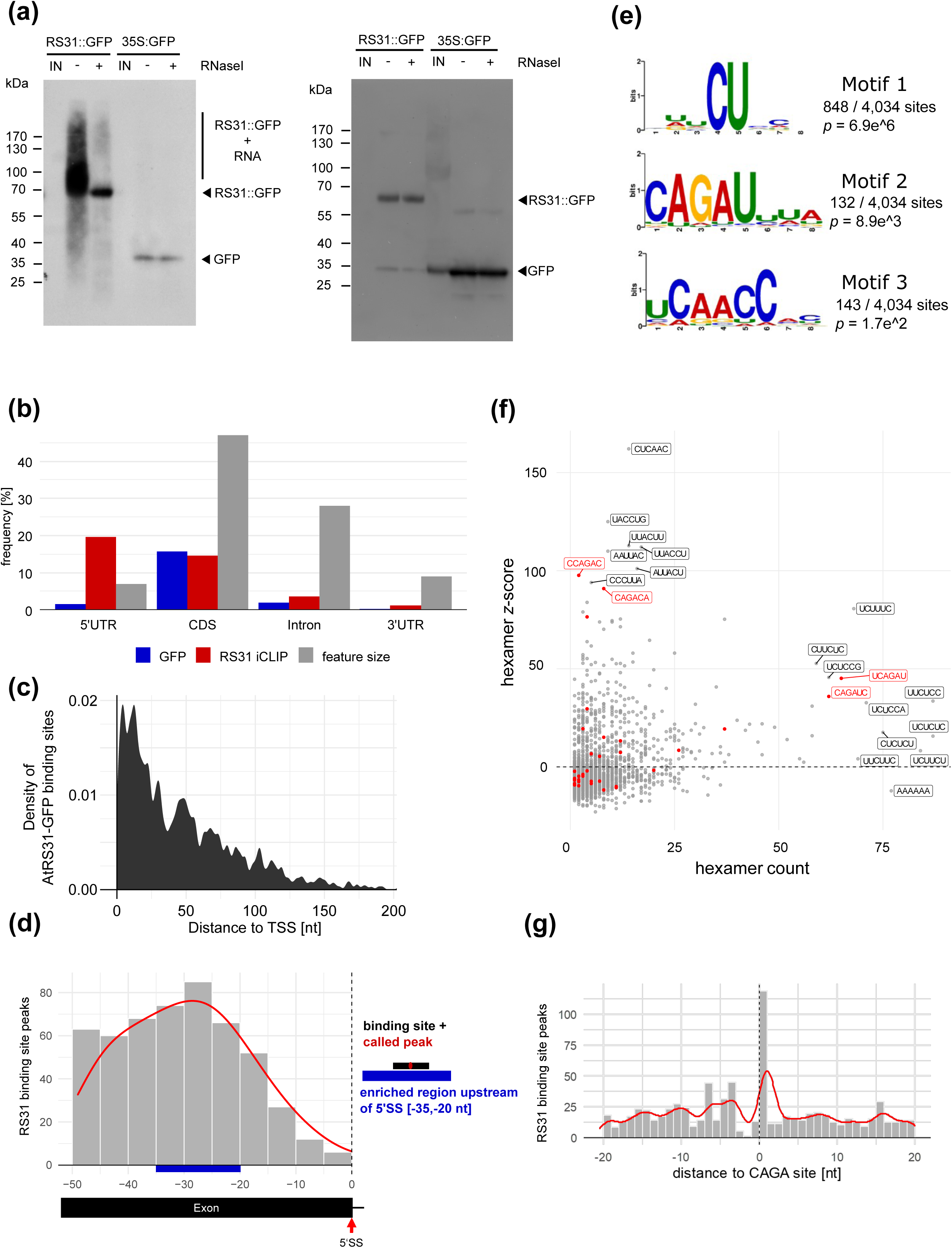
Determination of At-RS31 *in vivo* binding sites by iCLIP. (a) Left: Autoradiogram of RS31-GFP and GFP protein-RNA complexes. After UV crosslinking, cell lysates were subjected to immunoprecipitation with GFP Trap beads. RNAs were radioactively labelled, and the complexes were separated by denaturing gel electrophoresis. IN, input (lysate). Treatment of the precipitate with RNase I (+ RNase) indicates the size of the precipitated proteins. The region above the fusion protein containing the co-precipitated RNAs used for library preparation is indicated. Right: iCLIP western blot. Immunoblot analysis of the membrane shown in the left panel with anti-GFP antibody. Bands for GFP and RS31-GFP are marked accordingly. (b) Distribution of the At-RS31 binding sites within protein-coding transcripts. Distribution of RS31-GFP and GFP binding sites within transcripts in relation to the total length of the transcript features. 5’ UTR and 3’ UTR - 5’ and 3’ untranslated regions; CDS - coding sequences. (c) Distribution of the At-RS31 binding sites within 5’ UTRs relative to the transcription start sites (TSS). Distance to TSS is shown in nucleotides (nt). (d) Distribution of At-RS31 binding sites peaking upstream of 5’ splice sites (5’SS). Only exons at least 50 nucleotides in length were analyzed. The red line represents the local density of binding sites. The blue box marks the -35-20 nt region upstream of 5’SS where RS31-GFP binding sites are enriched. (e) Significant STREME binding site motifs. Sequence logos of the most significant (based on their p-value) RS31-GFP binding motifs identified by STREME analysis. For the analysis, only sequences from the 9-nucleotide binding site regions were considered. (f) Hexamer counts and z-scores. Scatterplot showing hexamer frequencies and counts computed from the 9-nucleotide binding site sequences of the RS31-GFP iCLIP sample. Hexamer counts (x-axis) are compared against hexamer z-scores (y-axis). The most enriched hexamers and the ones with highest counts are labelled according to their sequence. The highlighted hexamers (red) contain the subsequence CAGA. (g) iCLIP binding site distance to RNAcompete motif sites. Distribution of RS31-GFP binding site peaks in relation to CAGA sites across At-RS31 target transcripts. The position 1 on the x-axis denotes to the C in the CAGA motif identified by the RNAcompete assay.

PureCLIP was used to call peaks from iCLIP reads (Krakau *et al*., 2017). For RS31-GFP and GFP, 6,939 and 508 peaks were recovered, respectively (Table **S2**). If peaks were directly adjacent, we considered only the one with the highest score assigned by PureCLIP. Peaks were then extended by four nucleotides in both directions to define binding sites of nine nucleotides. This resulted in 4,236 binding sites for RS31-GFP and 324 binding sites for the GFP control. Of the 324 GFP only binding sites, 202 were located within nine nucleotides of RS31-GFP binding sites and were subtracted, leaving 4,034 RS31-GFP binding sites (Tables **S2** and **S3**) assigned to 1142 protein-coding and 279 non-coding genes (Table **S4**).

To gain insights into the At-RS31 targets and the molecular processes and biological pathways it may influence, we performed a functional enrichment analysis of the 1421 iCLIP target genes using g:Profiler (Kolberg *et al*., 2023) (Table **S5**). The analysis revealed a significant enrichment of genes involved in various stress responses, as well as in photosynthesis, carbon utilization and plant responses to light supporting our previous findings on the regulation of *At-RS31* alternative splicing by chloroplast retrograde signaling and TOR kinase in response to photosynthesized sugars and light (Petrillo *et al*., 2014; Riegler *et al*., 2021). Furthermore, At-RS31 targets are associated with RNA splicing, particularly mRNA splicing via the spliceosome, underscoring its role in co- and/or post-transcriptional regulation (Table **S5**).

Out of the 1142 protein-coding genes containing At-RS31 binding sites, 61.3% (700) have them in the 5’UTR, 43.2% (493) in the coding region (CDS), 11.7% (134) in introns, and 4% (46) in the 3’UTR (Fig. **1b**). Additionally, there is an accumulation of At-RS31 binding sites towards the transcriptional start site (TSS) in many genes (Fig. **1c** and Table **S6**), a pattern previously observed for putative *cis*-elements of the Arabidopsis SR-like protein SR45 (Xing *et al*., 2015). The functional significance of this distribution remains to be determined.

Since At-RS31 is a splicing regulator that interacts with the U1-70K and U11/U12-31K proteins, which are involved in the recognition of the 5’ splice site (Lorkovic *et al*., 2005; Altmann *et al*., 2020), we determined the position of the At-RS31 binding site peaks (center position of binding site) relative to 5’ splice sites (Fig. **1d**). For this, only exons with a length of at least 50 nt were considered. The distribution of binding site peaks has a slight enrichment at 25-30 nt upstream of the 5’ splice sites. Therefore, we asked whether these sites share conserved sequence motifs. Binding site peaks located 25-30 nt upstream of 5’ splice sites were extracted, extended to 21 nt and aligned (Fig. **1d**, Table **S7**). The alignment of the sequences consists mainly of C and U and is presented as a sequence logo in Fig. **S5**. Therefore, the polypyrimidine-rich sequences may impact the regulation of 5’ splice site choice by attracting At-RS31.

Investigating 4034 At-RS31 binding sequences using STREME revealed three enriched motifs (Fig. **1e**). The most significant pattern, motif 1, features a CU sequence, found at 848 binding sites (21 %). The second most significant motif, motif 2 (CAGAU), occurs in 132 sites (3.3 %), and motif 3, UCAACC, is present in 143 sites (3.5 %).

In parallel, we searched for hexamer motifs enriched within the binding sites. Among the motifs with the highest z-scored hexamers, predominantly CU and AC combinations are enriched (Fig. **1f**). CU combinations (polypyrimidines) appear among the most frequent but less enriched hexamers. In addition, the motif CAGA is present among the enriched and highest counts (highlighted in red, Fig. **1f**).

### Consistent and divergent RNA-binding motifs identified by RNAcompete and iCLIP for At-RS31

To validate the At-RS31 *in vivo* binding sites obtained through iCLIP, we employed RNAcompete that challenges RNA-binding proteins (RBPs) with a pool of short RNAs (30-41 nucleotides) that include almost every possible 7-mer sequence combination (Fig. **2a**) (Ray *et al*., 2013; Ray *et al*., 2017). For this, we generated constructs harboring full-length At-RS31 and its truncated version with the two RRMs but lacking the RS region, respectively (Fig. **S2a**). The corresponding GST fusion proteins, GST-31 FL and GST-31 RRMs, were purified from *E. coli* and their integrity was verified (Fig. **S2b**).

**Fig. 2.**
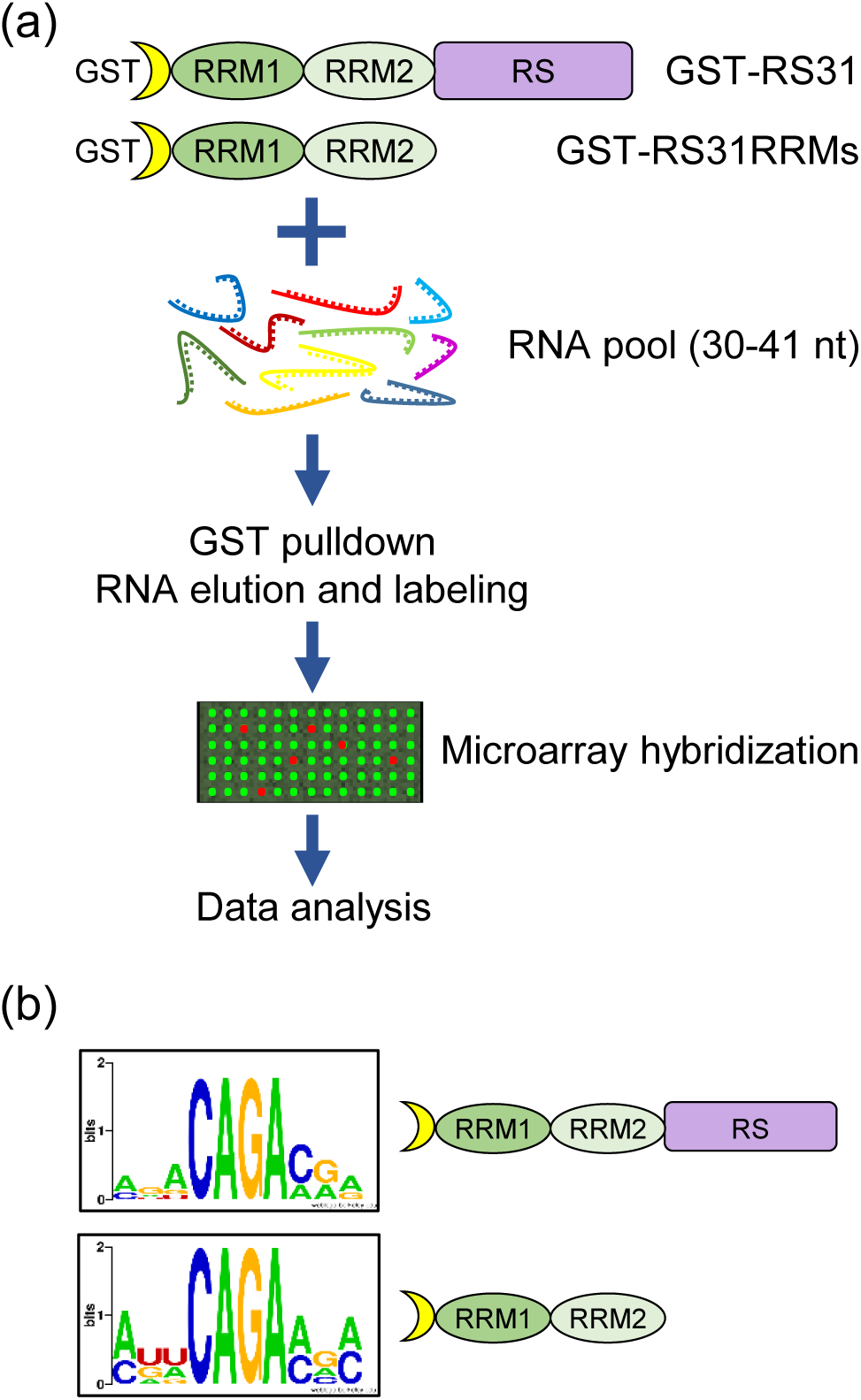
Identification of At-RS31 RNA-binding motifs using RNAcompete. (a) Overview of the RNAcompete assay. GST-tagged full-length At-RS31 protein and its truncated version comprising both RRMs were incubated with a 75-fold molar excess of designed RNA pool. RNA bound to GST-RS31 and GST-RS31RRMs fusion proteins during the GST pulldown was eluted, purified, labeled and hybridized to custom Agilent 244K microarrays. Microarray data was analysed computationally to identify 7-mers specifically bound by At-RS31 and generate RNA-binding motifs. RRM1 and RRM2 – RNA recognition motif domains; RS – region rich in arginines and serines. (b) RNA-binding motifs of the full-length At-RS31 and its truncated version containing RRMs identified in the RNAcompete assay and represented as sequence logos.

The RNAcompete experiments revealed AGACAGA as the highest-scoring 7-mer (Fig. **S6**). Intriguingly, the top ten motifs bound by GST-31 FL and GST-31 RRMs contain a core 4-mer CAGA (Fig. **2b** and Fig. **S6**), suggesting that both variants have very similar *in vitro* binding properties, and that the RS region does not contribute significantly to At-RS31 binding specificity *in vitro*. The CAGA 4-mer was also present at the At-RS31 binding sites identified by iCLIP (motif 2, Fig. **1e**) and within high-scoring hexamers (Fig. **1f**).

We further analyzed the distance of the At-RS31 iCLIP binding site peaks relative to the CAGA motif (Fig. **1g**). The results revealed a positive correlation between the binding sites and the CAGA motif, with a higher number of binding site peaks positioned towards the cytosine (C) of the CAGA motif.

Overall, the RNA-binding motifs identified by RNAcompete for At-RS31 are highly consistent with the motifs identified by iCLIP, although iCLIP has identified additional motifs (see Fig. **1e**). This difference may be attributed to the fact that iCLIP also reflects post-translational modifications and protein-protein interactions of At-RS31 *in planta*. Hence, the combination of both *in vitro* and *in vivo* methods provides a more comprehensive understanding of the RNA-binding preferences of At-RS31 and highlights the importance of considering the context in which RBPs proteins function.

### At-RS31 shapes alternative splicing patterns and gene expression in Arabidopsis

To better understand the biological processes influenced by At-RS31 and its role in regulating gene expression, we performed a transcriptome analysis using RNA-seq on the *rs31-1* mutant, plants overexpressing the mRNA1 protein-coding isoform (*35S::RS31*), and wild-type (wt) control plants (Fig. **3b-d**). Under normal growth conditions, *35S::RS31* and *rs31-1* plants show only minor phenotypic differences wt plants, with *35S::RS31* plants being slightly smaller (Fig. **3d**) (Petrillo *et al*., 2014).

**Fig. 3.**
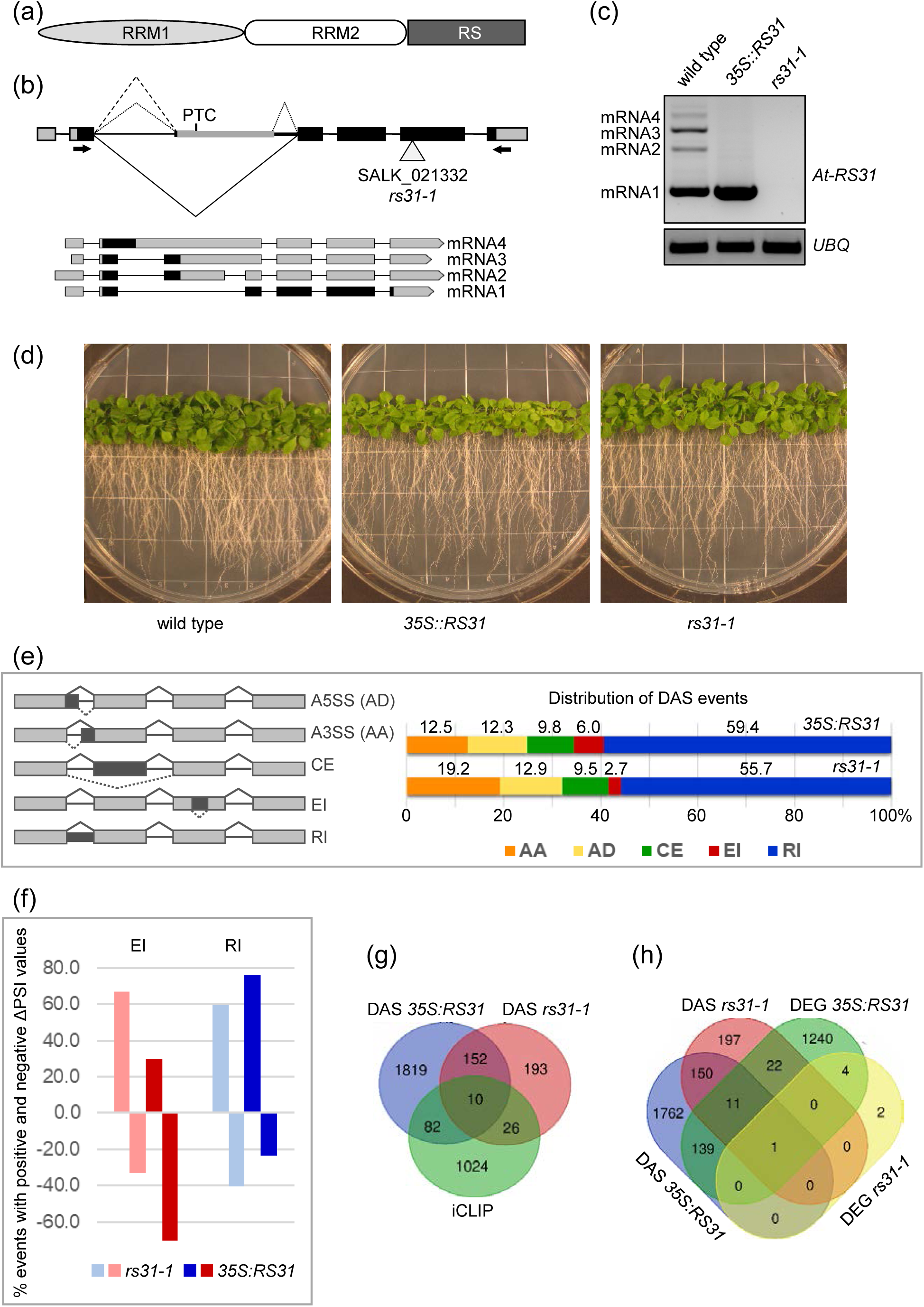
At-RS31 and its impact on alternative splicing and gene expression in *Arabidopsis thaliana*. (a) Schematic of the At-RS31 protein domain structure. The protein contains two RNA recognition motifs (RRM1 and RRM2) and an arginine/serine-rich region (RS). (b) Structure of At-*RS31* gene, its alternative splicing events and transcript models. Carets indicate alternative splicing events in the second intron. The solid caret represents the splicing event that produces the reference transcript AT3G61860.1 (mRNA1), which encodes the At-RS31 protein. Retention of the second intron generates the mRNA4 transcript. The dashed caret marks the use of an alternative 3’ splice site, resulting in the mRNA3 transcript. Dotted carets denote an alternative exon generated via use of both alternative 3’ and 5’ splice sites, producing the mRNA2 variant. PTC - the premature termination codon. Regions from the AUG start codon to the reference stop codon (mRNA1) or the PTC (mRNA2-4) are shaded in black. The arrowhead indicates the T-DNA insertion site in the SALK_021332 line (*rs31-1* mutant). Black arrows show positions of primers used for RT-PCR analysis in (C). (c) RT-PCR analysis of At-*RS31* transcript levels in the *A. thaliana* wild type, *35S::RS31* (overexpression of the mRNA1 under the CaMV 35S promoter), and *rs31-1* plants. Ubiquitin (*UBQ*) was used as a loading control. (d) Phenotypes of wild type, *35S::RS31* overexpression, and *rs31-1* mutant plants used for RNA-sequencing. Plants were grown under a 16 h light/8 h dark cycle at 22°C on vertical agar plates containing half-strength MS medium. (e) Distribution of alternative splicing event types differentially regulated in *rs31-1* and *35S::RS31* lines in comparison to wild type *A. thaliana*. Diagrams on the left illustrate the analyzed alternative splicing event types: AA (alternative acceptor, or alternative 3’ splice site), AD (alternative donor, or alternative 5’ splice site), CE (cassette exon), EI (exitron), and RI (retained intron). (f) Proportions of differential exitron (EI) and retained intron (RI) events with reduced or increased splicing efficiency in *rs31-1* and *35S::RS31* lines compared to wild type. Positive and negative percentages indicate the proportion of events with increased or decreased Percent Spliced In values (ΔPSI), respectively. (g) Comparison of differential alternative splicing (DAS) and iCLIP statistics. Venn diagram shows the overlap between genes with At-RS31 iCLIP binding sites and DAS genes in *rs31-1* and *35S::RS31* lines compared to wild type. (h) Comparison of DAS and DEG statistics. Venn diagram shows numbers of genes differentially alternatively spliced (DAS) and differentially expressed (DEG) in *rs31-1* and *35S::RS31* lines compared to wild type.

We identified 442 differential alternative splicing (DAS) events across 381 genes in *rs31-1* plants, and 2949 DAS events across 2063 genes in *35S::RS31* plants, using a change in Percent Spliced In value |ΔPSI|≥ 0.1 as a threshold (Table **S8**). Interestingly, 42.5% (162 out of 381) of the genes affected by DAS in *rs31-1* plants were also affected in *35S::RS31* plants. However, only ∼30% (130 out of 442) of the specific DAS events were shared between the two lines (Table **S8**), suggesting that alterations in At-RS31 levels lead to distinct splicing outcomes within the same genes, depending on whether it is underexpressed or overexpressed.

Previous studies in plants have consistently reported that alternative 3′ splice sites (alternative acceptor, AA) are generally used about twice as frequently as alternative 5′ splice sites (alternative donor, AD) (Wang & Brendel, 2006; Marquez *et al*., 2012; Zhang *et al*., 2022). This ratio is nearly constant across species in different kingdoms (McGuire *et al*., 2008). However, in *rs31-1* and *35S::RS31* plants, the AA/AD ratio was reduced to ∼1.4:1 and ∼1:1, respectively (Fig. **3e**, Table **S8**), suggesting that At-RS31 may influence this balance. One explanation could be that At-RS31 modulates the recognition and usage of a subset of 5’ splice sites, as implied by the distribution of binding site peaks 25-30 nt upstream of 5’ splice sites (Fig. **1d**).

Intron retention (RI) was the most frequent DAS event type in both *rs31-1* (55.7%) and *35S::RS31* (59.4%) plants (Fig. **3e**), a higher prevalence than the ∼40% reported for wt plants (Marquez *et al*., 2012). We have previously shown that RIs and exitrons (EIs), defined as alternatively spliced internal regions of protein-coding exons, have distinguishable features, including their regulation and functional outcomes (Marquez *et al*., 2015; Jabre *et al*., 2021). Indeed, the direction of splicing changes for RIs and EIs was significantly different (p-value 7.51×10^-39^) in *35S::RS31* plants, where retention of introns was predominantly promoted (76.2%), while splicing of EIs was predominantly enhanced (70.2%) (Fig. **3f**). Conversely, in *rs31-1* plants, splicing of EI was inhibited in 66.7% of DAS EIs (Table **S8**). These contrasting patterns of EI splicing between *rs31-1* and *35S::RS31* plants suggest a positive role for At-RS31 in promoting EI splicing.

The comparison between DAS and iCLIP data sets revealed that At-RS31 binds to the transcripts of 36 and 92 DAS genes in *rs31-1* and *35S::RS31,* respectively (Fig. **3g**, Table **S8**). DAS events in selected genes with At-RS31 binding sites were validated using RT-PCR (Fig. **S7**). GO term analysis showed that DAS genes with At-RS31 iCLIP binding sites were associated with terms related to mRNA splicing via spliceosome, spliceosomal complex, and nuclear speck (Table **S5**). This further underscores role of At-RS31 in alternative splicing regulation and its integration into the splicing machinery.

The impact of At-RS31 on alternative splicing was more pronounced than its effect on gene expression (Fig. **3h**). Overexpression of At-RS31 resulted in the differential expression (DE) of 1417 genes, with 798 up- and 619 down-regulated genes (ILog_2_FCI > 1, Table **S9**). Only six DEGs were identified in the *rs31-1* mutant, excluding *At-RS31* itself. Notably, 7.3% (151 out of 2063) of *35S::RS31* DAS genes were also differentially expressed (Fig. **3h**), indicating largely separate effects of At-RS31 on alternative splicing and gene expression.

Cross-referencing the DE and iCLIP data showed that At-RS31 binds to the transcripts of 77 *35S::RS31* DEGs (Table **S9**). The balanced distribution between upregulated (41) and downregulated (36) genes with At-RS31 binding sites suggests that At-RS31 can have both positive and negative effects on gene expression. However, it is possible that much of the differential expression observed may be downstream consequences of At-RS31 direct regulatory function on splicing.

The impact of At-RS31 on transcription factor (TF) splicing may account for some of the observed differential expression *via* alternative TF isoforms with altered regulatory properties. Out of 2,534 TFs (Calixto *et al*., 2018), At-RS31 binding sites were found on the transcripts of 107 TFs, while 192 and 39 TFs showed DAS in *35S::RS31* and *rs31-1* plants, respectively (Tables **S8**, TF column, and **S10**). For example, At-RS31 binds to *WRI4/AP2-2* (AT1G79700), which responds to abscisic acid (ABA), light and carbohydrate availability (Vogel *et al*., 2012). It promotes two in-frame AS events (AD and CE removing 10 and 47 amino acids, respectively) that affect the AP2 DNA-binding domain and may influence its regulatory function. These splicing changes may influence TF regulatory functions and lead to alterations in the expression of their target genes.

Finally, GO term enrichment of DEGs in the *35S::RS31* plants revealed a strong association with responses to stress, including hypoxia, chemical stimuli, and responses to both abiotic and biotic stimuli (Table **S5**). This contrasts the enrichment patterns observed in DAS gene sets, corroborating the small overlap between DAS and DE genes (Fig. **3h**) and emphasizing the separate mechanisms by which At-RS31 influences alternative splicing and gene expression.

### At-RS31 has a broad impact on RNA-binding proteins and splicing factors

GO term analysis revealed an enrichment in splicing and mRNA processing terms in the At-RS31 iCLIP and DAS gene datasets (Table **S5**). Moreover, the low overlap between iCLIP peaks and At-RS31-affected genes suggested that the regulatory impact of At-RS31 could partly be indirect, likely by modulating other splicing factors (SFs). To investigate this, we examined the effects of At-RS31 on alternative splicing of 798 known SFs and RBPs (Calixto *et al*., 2018). 184 DAS events were identified in 117 SF/RBPs in *35S::RS31* plants, and 27 DAS events were found in 17 genes in *rs31-1* plants (Table **S8**, SF-RBP column). In contrast, differential expression was observed in far fewer SF/RBPs, with 13 DEGs in *35S::RS31* and none in *rs31-1* (Table **S9**, SF-RBP column).

RT-PCR validation confirmed the DAS events in selected SF and RBPs (Fig. **S8**). Interestingly, At-RS31 regulates EI splicing in some of these genes. For example, At-RS31 inhibits splicing of an in-frame EI in the flowering regulator *FPA*, which controls mRNA 3′ end formation (Hornyik *et al*., 2010), thereby favouring production of the full-length protein (Fig. **S8**, Table **S8**). Similarly, the inhibition of out-of-frame EI splicing results in the production of full-length proteins, as observed in the snRNA transcription activator *SRD2* (Ohtani & Sugiyama, 2005), the splicing factor *RDM16,* required for RNA-directed DNA methylation (Huang *et al*., 2013), the mRNA export factor *SAC3a* (Kanno *et al*., 2018), and the cold-regulated kinase *CDKG1* (Cavallari *et al*., 2018).

Additionally, At-RS31 binding sites were identified in 68 SF and RBPs (Table **S4**). Of these, 17 underwent DAS in *35S::RS31* plants, and three were affected in both *rs31-1* and *35S::RS31* (Table **S8**). Besides SR proteins, this group includes CCR1/At-GRP8, an hnRNP-like glycine-rich RNA-binding protein (Streitner *et al*., 2012), CypRS64, a cyclophilin that interacts with SR proteins and components of U1 and U11 snRNPs (Lorkovic *et al*., 2004), and CIS1, a splicing factor involved in blue-light-mediated flowering via interaction with cryptochrome CRY2 (Zhao *et al*., 2022). The presence of At-RS31 binding sites in these and other SF/RBPs suggests that they are likely direct targets of At-RS31, positioning them as key players in its regulatory network.

### Cross-regulation of SR protein family by At-RS31

SR proteins are regulated by alternative splicing, which involves both auto-regulation and cross-regulation within the SR protein family (Lopato *et al*., 1999; Sureau *et al*., 2001; Kalyna *et al*., 2003; Kalyna *et al*., 2006; Palusa *et al*., 2007; Müller-McNicoll *et al*., 2019). Here, we identified that two paralogs of *At-RS31, At-RS31a* and *At-RS40*, along with a member of the ASF/SF2 subfamily, *At-*SR34a (Barta *et al*., 2010), display differential alternative splicing in the *rs31-1* mutant. Additionally, in the *35S::RS31* overexpression line, 12 of the 18 Arabidopsis SR genes demonstrate DAS, with *At-RS41* also showing differential expression (Fig. **4a**, Tables **S8** and **S9**). iCLIP analysis revealed At-RS31 binding peaks in *At-RS31a, At-RS40*, *At-RS41, At-SR30, At-SCL30a,* and *At-SCL33,* suggesting their regulatory significance (Fig. **4**, Tables **S4** and **S8**). Additionally, 13 SR genes contain top 7-mers identified through RNAcompete (Fig. **4a**).

**Fig. 4.**
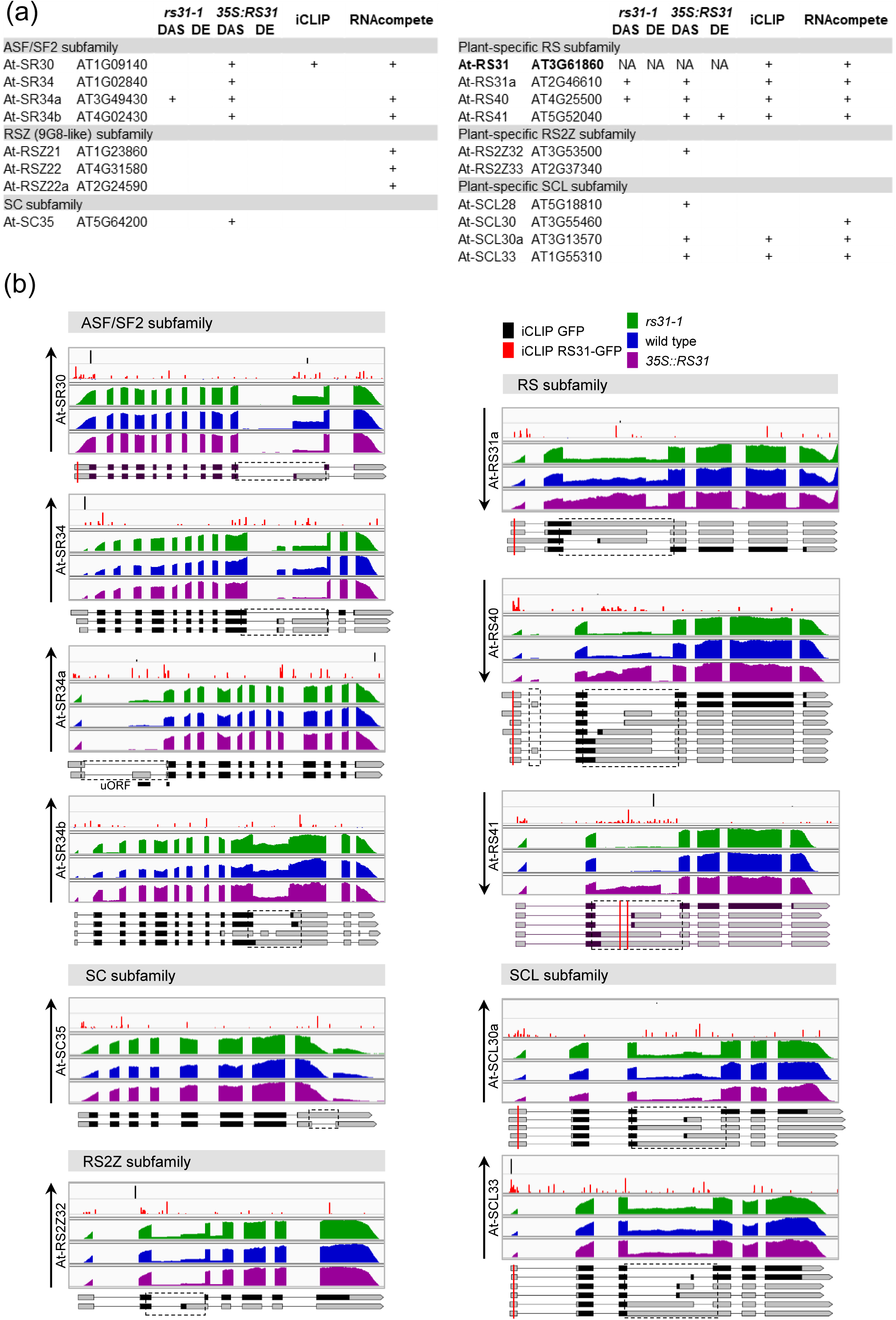
Regulation of Arabidopsis SR protein family by At-RS31. (a) Overview of differential alternative splicing (DAS) and differential gene expression (DE) in Arabidopsis SR genes in *rs31-1* mutant and *35S::RS31* overexpression plants, compared to wild-type controls. The table includes information on iCLIP peaks and the top 7-mers identified by RNAcompete within SR genes. “NA” indicates data not applicable. (b) Integrated Genomics Viewer (IGV) tracks depicting iCLIP cross-link sites for GFP and RS31-GFP, along with RNA-seq read coverage tracks for *rs31-1*, *35S::RS31*, and wild-type plants. Transcript models for SR genes are displayed using the Boxify tool (https://boxify.boku.ac.at/). Protein-coding regions, spanning from the translational start codon to the stop codon or premature termination codon, are shown in black. Red vertical lines represent the locations of At-RS31 binding sites identified by iCLIP. Only transcripts relevant to the identified DAS events are displayed, with dashed rectangles marking these alternatively spliced regions. A black line denotes the position of an upstream open reading frame (uORF) in *At-SR34a*. Black upward and downward arrows indicate an increase or decrease, respectively, in the ratio of splice variants encoding full-length SR proteins.

All ASF/SF2 subfamily members (*At-SR30, At-SR34, At-SR34a,* and *At-SR34b*) display DAS events influenced by At-RS31 (Fig. **4**, Table **S8**). A notable example is CE skipping in the 5’ UTR of *At-SR34a*, promoted in *35S::RS31* plants but inhibited in *rs31-1* mutant. CE inclusion creates an upstream open reading frame (uORF), overlapping the primary ORF start site, leading the transcript to NMD (Kalyna *et al*., 2012). Thus, At-RS31 likely upregulates At-SR34a by controlling AS to counteract NMD. Overexpression of At-RS31 also favours the protein-coding isoforms of *At-SR30, At-SR34,* and *At-SR34b* (Fig. **4b**, Table **S8**). Therefore, At-RS31 regulates AS in this subfamily, enhancing protein-coding isoforms without changing overall gene expression.

In the RS2Z, SCL, and SC subfamilies, At-RS31 also reduces unproductive AS isoforms, thus increasing the proportion of protein-coding isoforms (Fig. **4b**, Table **S8**). RS2Z and SCL genes contain remarkably long introns located between RNP2 and RNP1 motifs of their N-terminal RRMs. These introns undergo highly conserved AS from the moss *Physcomitrium patens* to Arabidopsis, suggesting their critical regulatory roles (Kalyna & Barta, 2004; Iida & Go, 2006; Kalyna *et al*., 2006). For example, in *At-RS2Z32*, At-RS31 inhibits the usage of a conserved alternative 3’splice site (Fig. **4b**, Table **S8**). Similar to the AS events regulated by At-RS31 in the ASF/SF2 genes, the DAS events in *At-RS2Z32, At-SCL30a*, and *At-SCL33* (Fig. **4b** and Table **S8**) predominantly result in AS transcripts sensitive to NMD (Palusa & Reddy, 2010; Kalyna *et al*., 2012). Interestingly, in *At-SC35*, At-RS31 inhibits splicing of an intron within the 3’ UTR, preventing NMD and promoting the accumulation of the protein-coding variant (Palusa & Reddy, 2010) (Fig. **4b**, Table **S8**). A similar regulatory mechanism was found in the overexpression of SC35 in human cells (Sureau *et al*., 2001).

Conversely, in the plant-specific RS subfamily, which includes At-RS31 and its paralogs At-RS31a, At-RS40, and At-RS41, At-RS31 favours unproductive AS variants, reducing the proportion of protein-coding isoforms (Figs. **4b** and **S8**, Table **S8**). The AS in these genes also occurs in their longest introns and is conserved from Arabidopsis to green alga *Chlamydomonas* (Kalyna *et al*., 2006). iCLIP analysis shows accumulation RS31-GFP peaks within these introns, suggesting that At-RS31 directly interacts with these regions (Fig. **4b**). *At-RS31a* and *At-RS40* show opposite splicing regulation in *rs31-1* and *35S::RS31* contexts. At-RS31 promotes a shift toward CE-containing NMD-sensitive transcripts (Kalyna *et al*., 2012; Fuchs *et al*., 2021), decreasing the proportion of protein-coding transcripts. *At-RS41* follows a similar pattern but only in *35S::RS31* plants (Figs. **4b** and **S8**, Table **S8**). Overall expression of *At-RS40* and *At-RS41* is reduced in *35S::RS31* plants (Table **S8**). This demonstrates negative regulation by At-RS31 of its paralogs through AS coupled to NMD, reducing the amount of their functional proteins.

Taken altogether, At-RS31 increases the abundance of protein-coding isoforms for most SR genes, except for its paralogs, where it promotes the accumulation of non-productive transcripts. At-RS31 serves as a general regulator of protein-coding isoforms across different subfamilies of SR proteins in Arabidopsis, acting as a hierarchical regulator of alternative splicing.

### At-RS31 and TOR pathway share targets regulating key aspects of plant growth and stress response

The TOR signalling pathway influences cell growth and metabolism in response to nutrients, growth factors, and environmental signals (Burkart & Brandizzi, 2021). Our previous research showed that TOR kinase regulates alternative splicing of *At-RS31* in response to sugars and light, increasing the proportion of the protein-coding splice variant (Riegler *et al*., 2021). To determine if At-RS31 target genes may be regulated through the crosstalk with the TOR pathway, we analysed 160 genes undergoing DAS in response to TOR activity modulation by glucose and the TOR inhibitor Torin2 (Riegler *et al*., 2021) (Table **S11**). Of these, 7 (4.2%, p-value 0.0024) and 43 (27%, p-value 8.40×10^−17^) showed DAS in the *rs31-1* and *35S::RS31* lines, respectively. Additionally, 13 (8.1%, p-value 0.0033) of these genes have At-RS31 binding sites identified by iCLIP, with six of these showing DAS in the At-RS31 overexpression plants, suggesting they could be direct targets for TOR-mediated regulation of alternative splicing via At-RS31.

Among these putative targets, CIPK3, a kinase regulating abscisic acid (ABA) stress responses and seed development (Kim *et al*., 2003), has an At-RS31 binding site 20 nt upstream of an unannotated CE containing a PTC (Table **S8**, Fig. **S9a**). At-RS31 promotes exclusion of this exon, potentially increasing the proportion of the full-length protein. Notably, CIPK3 interacts with RAPTOR1B and may negatively regulate TORC activity by phosphorylation (Li *et al*., 2023), suggesting a link between At-RS31-mediated splicing and TOR activity, particularly under ABA-dependent conditions.

Further analysis of At-RS31 iCLIP targets and DAS genes (Tables **S4** and **S8**) revealed additional TOR targets not present in the initial dataset (Riegler *et al*., 2021). The *MCM3* gene (AT5G46280), DNA replication helicase and a known TOR marker gene (Yamamoto *et al*., 2018; Jamsheer K *et al*., 2022b), is negatively regulated by At-RS31. In *rs31-1* mutant, *MCM3* intron retention decreases, while it increases in *35S::RS31* plants (Table **S8**, Fig. **S9b**), implying a role for alternative splicing in regulating MCM3 protein levels. At-RS31 binding sites were identified on transcripts of S6K1 kinase and RPS6B ribosomal protein (Table **S4**), key components of the TOR pathway and TOR phosphorylation targets that modulate translational capacity of the cell (Dobrenel *et al*., 2016). This raises the possibility that At-RS31 influences alternative splicing of TOR phosphoproteins, adding a regulatory layer upstream of TOR.

To investigate this further, we examined 160 genes encoding TOR-phosphorylated proteins (Van Leene *et al*., 2019; Scarpin *et al*., 2020) (Table **S11**). At-RS31 binding sites were found on the transcripts of 18 of these genes (p-value 9.28×10^−6^), and 16 exhibited DAS (p-value 0.0373). Five genes (*ATG13, NIG, P2Y/RPP2B, At-RS40*, and *At-RS41*) showed both At-RS31 binding and DAS, likely representing direct targets of At-RS31. The shared genes are involved in essential processes such as metabolism, autophagy, cytoskeleton organization, and RNA splicing (Table **S11**).

We also investigated proteins interacting with TORC1 components that represent genuine TOR targets and upstream regulators (Jamsheer K *et al*., 2022a). Among 167 TORC1 interactors, At-RS31 binding sites were identified in 13 genes (p-value 0.0048), including the TOR target S6K1, while 18 showed DAS (p-value 0.0144) (Table **S11**). Several in-frame DAS events have the potential to generate protein isoforms (Table **S11**). They include exitrons in pyruvate dehydrogenase EMB3003, tetratricopeptide repeat-like protein TPR1, and eukaryotic translational initiation factor eIF2B-ε3. In the deubiquitinase UBP12, DAS leads to an isoform differing by one amino acid due to an alternative 3’ splice site. In TTI1, part of the Triple T complex, At-RS31 modulates an alternative 5’ splice site, leading to inclusion of six amino acids, possibly influencing TOR activation in response to glucose/energy status of the cell (Liu & Xiong, 2022). In some genes, DAS occurs in UTRs, whose impact is not always clear (Table **S11**). However, At-RS31 may modulate presence of uORFs, as observed for the ABA-activated SnRK2.8 kinase (Wu & Hsu, 2023), where At-RS31 inhibits intron retention in the 5’ UTR, potentially removing a uORF that may enhance translation of the main ORF. Multiple At-RS31 iCLIP crosslinks in the SnRK2.8 5’ UTR suggest a direct regulatory role (Tables **S8** and **S11**, Fig. **S9c**). Given that SnRK2.8 interacts with Raptor1B, whose phosphorylation by SnRK2s disrupts the TOR complex to limit growth under stress (Wang *et al*., 2018), At-RS31 may act upstream, controlling ABA-dependent TOR inactivation. Additionally, At-RS31 binding site is present in the ABA receptor PYL4, which also interacts with TOR (Table **S11**) (Wang *et al*., 2018). Together, several examples suggest a broader role for At-RS31 in coordinating responses between TOR-mediated growth control and ABA signalling.

The overlap of genes affected by both TOR modulation and At-RS31 activity reveals a complex, multi-level regulatory control. These genes appear to be controlled post-transcriptionally via At-RS31-mediated splicing and possibly at the translational level via TOR-dependent mechanisms.

### Role of At-RS31 in integrating abscisic acid metabolism and signaling with the TOR pathway

We observed that among genes that undergo DAS upon TOR inhibition or encode TOR-dependent phosphoproteins and TORC1 interactors (Table **S11**), At-RS31 binds and modulates the alternative splicing of the transcripts of several important genes related to the ABA pathway. Moreover, functional enrichment analysis of At-RS31 targets revealed GO terms associated with response to ABA and other stresses (Table **S5**). Increased ABA levels inhibit TOR signalling, thereby restricting growth under stress, while TOR counteracts ABA signalling to promote growth under favourable conditions. This interplay balances plant growth and stress responses (Wang *et al*., 2018). Given that At-RS31 is regulated by TOR (Riegler et al., 2021) and influences alternative splicing of TOR-related genes (Table **S11**), including those involved in the ABA pathway, we hypothesized that At-RS31 serves as a molecular link between the TOR and ABA pathways. To test this hypothesis, we examined the effects of At-RS31 overexpression and knockout on ABA responses in Arabidopsis seedlings. While *rs31-1* seedlings exhibited responses similar to wild-type plants, At-RS31 overexpression significantly enhanced ABA sensitivity, resulting in marked growth inhibition and a paler appearance (Fig. **5a**).

**Fig. 5.**
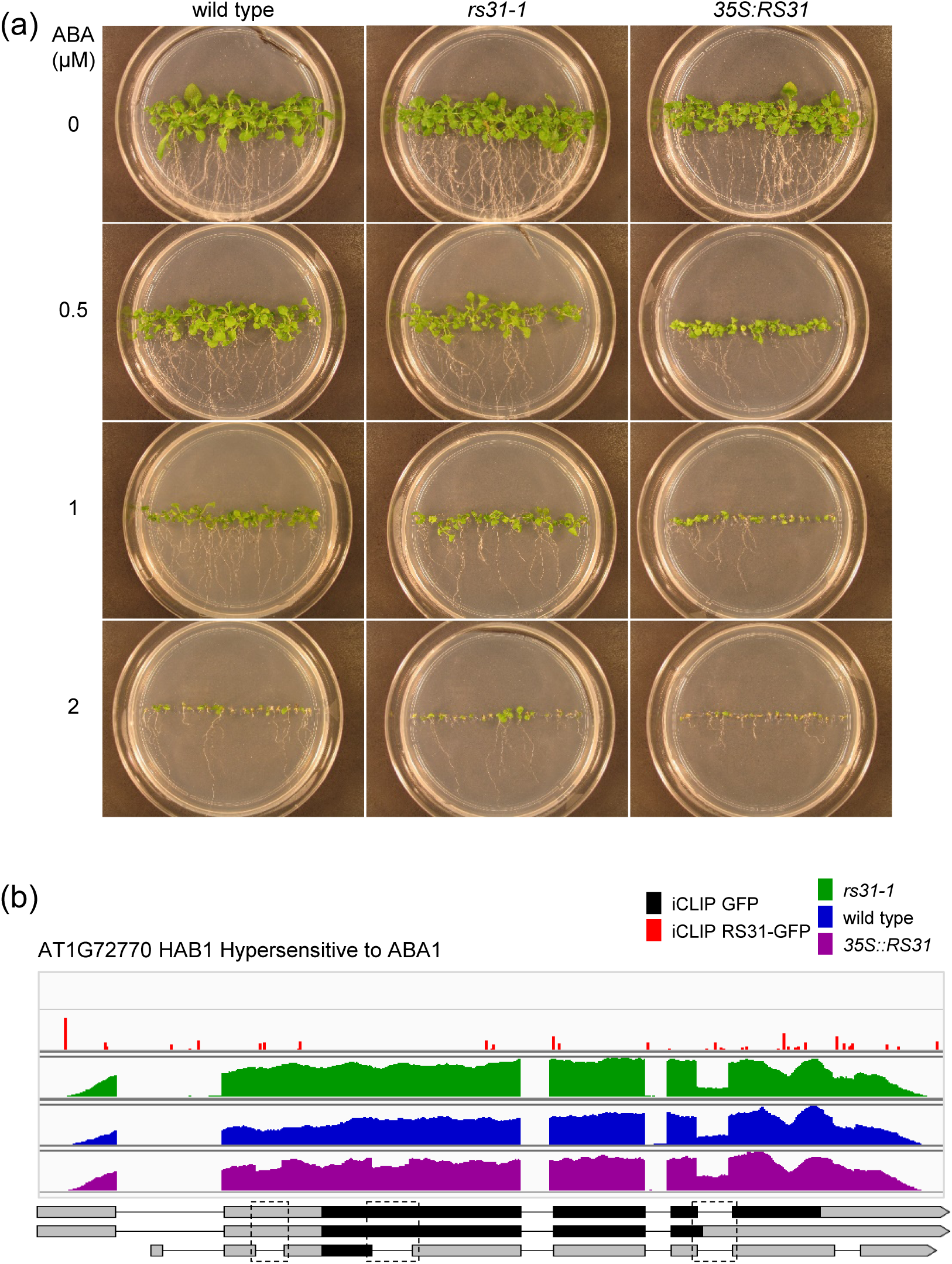
At-RS31 is involved in regulating abscisic acid pathway. (a) Phenotype of At-RS31 mutant and overexpression plants in response to abscisic acid. Seeds of *rs31-1* mutant, *35S::RS31* overexpression, and wild-type plants were germinated and grown on the vertical MS agar plates supplemented with 0, 0.5, 1, and 2 µM of abscisic acid at 23°C, 16h light/8h dark. (b) At-RS31 modulates alternative splicing of HAB1 (Hypersensitive to ABA1). Integrated Genomics Viewer (IGV) tracks show iCLIP cross-link sites for GFP and RS31-GFP, along with RNA-seq read coverage for *rs31-1*, *35S::RS31*, and wild-type plants. In the transcript models, protein-coding regions, spanning from the translational start codon to the stop codon or premature termination codon, are shown in black. Only transcripts relevant to the identified DAS events are displayed, with dashed rectangles marking these alternatively spliced regions.

To determine whether this phenotype is linked to the regulation of ABA-related genes by At-RS31, we analysed its potential influence on a compiled set of 313 genes (Finkelstein, 2013), covering genes involved in ABA metabolism, transport, signalling, and core ABA-induced and ABA-repressed genes. In total, 89 out of these 313 genes contained At-RS31 binding sites or showed DAS or DE changes (p-value 8.49×10^-11^) (Table **S12**). Among these, 49 had At-RS31 binding sites identified by iCLIP (p-value 1.72×10^−14^), including genes involved in ABA synthesis (*NCED6*), catabolism (*CYP707A3*), conjugation (*UGT73B1*), and transport (*ABCG25*). At-RS31 also binds transcripts of ABA receptors PYL4 and PYL5, as well as GUN5/ABAR/CHLH, which plays a role in chloroplast retrograde signalling and antagonizes the WRKY18/40/60 transcription repressors of ABA signalling. Additionally, transcripts of several key transcription factors, phosphatases and kinases involved in ABA metabolism and signalling (*e.g*. WRKY18, WRKY40, AHG3, FRY2/CPL1, CRK36, CIPK3, and CPK32) were identified as At-RS31 targets. Notably, At-RS31 binding sites were more prevalent in core ABA-repressed genes (18 out of 47) than in ABA-induced genes (2 out of 68) (Finkelstein, 2013) (Table **S12**). Moreover, At-RS31 upregulates core ABA-repressed genes while down-regulating ABA-induced genes. In total, 28 ABA-related genes were present among *35S::RS31* DEGs (p-value 1.66×10^-4^) (Tables **S9** and **S12**).

At-RS31 also influences alternative splicing in ABA-related genes, though the number of affected genes is relatively low (3 genes in *rs31-1* and 22 in *35S::RS31*) (Table **S12**). Nevertheless, even this limited number of modulated splicing events could have meaningful impacts on the ABA pathway. The affected genes span various functional categories critical to ABA signalling and function, including ABA synthesis (*ABA3*), transport (*ABCG40/PDR12*), transcription regulation (e.g. *EEL* and *ABF3*), and ABA signalling components (Tables **S8** and **S12**).

Beyond DAS events in *CIPK3* and *SnRK2.8* (Fig. **S9**) described above, HAB1 represents another significant example of an ABA signalling component influenced by At-RS31. HAB1, a PP2C phosphatase, is a negative regulator of ABA signaling that dephosphorylates and inactivates SnRK2 kinases (Wang *et al*., 2018). We detected three DAS events in *HAB1* (Table **S8**, Fig. **5b**). At-RS31 promotes the removal of an intron within the 5’ UTR, and also the removal of an exitron in the first coding exon, introducing a PTC. These novel DAS events reduce the *HAB1.1* splice variant, which encodes the full-length protein. Importantly, At-RS31 also promotes retention of an intron between the third and fourth coding exons, generating the *HAB1.2* isoform, known to upregulate ABA signalling by acting antagonistically to *HAB1.1* (Wang *et al*., 2015).

Taken together, the regulation of *SnRK2.8* and *CIPK3* by At-RS31, both of which interact with the TOR complex and may inhibit its activity, is consistent with the need for enhanced ABA signalling under stress conditions. Additionally, modulation of the *HAB1* splicing by At-RS31 to favour *HAB1.2* isoform, which enhances ABA signalling, further supports its role in balancing ABA and TOR pathways. At-RS31 modulates expression of a wide range of ABA-related genes. By promoting alternative splicing events that enhance or repress specific ABA pathway components, At-RS31 provides an additional regulatory layer that integrates ABA and TOR pathways. This integration ensures that under stress conditions, ABA signalling is augmented to inhibit the TOR pathway, prioritizing stress responses over growth. Conversely, under favourable conditions, TOR signalling can regulate At-RS31 to support growth by modulating ABA signalling components.

## DISCUSSION

At-RS31, a plant-specific SR protein characterized by two N-terminal RRMs and a uniquely structured RS region (Fig. **3A**) (Lopato *et al*., 1996), demonstrates intriguing RNA-binding specificities. The RRM1 of At-RS31, containing the highly conserved RDAEDA region, shares 40-48% identity with the RRM1 domains of human group 1 SR proteins, specifically SFRS1, SFRS4, SFRS5, SFRS6 (Lopato *et al*., 1996; Manley & Krainer, 2010). This identity level is just below the 50% threshold typically indicative of similar RNA-binding preferences (Ray *et al*., 2013). Using *in vivo* iCLIP and *in vitro* RNAcompete, we identified consistent motifs that At-RS31 binds to in target transcripts, underscoring specific interaction of At-RS31 with its target transcripts. iCLIP revealed an enrichment of the CAGA motif in At-RS31 binding sites while RNAcompete confirmed binding preferences for CAGA-containing sequences, validating these *in vivo* targets. Interestingly, iCLIP also uncovered additional motifs, such as CU-rich motifs predominantly enriched upstream of 5’ splice sites, which were not detected by RNAcompete. Similarities and differences found using these approaches suggest that this SR protein may bind RNA in a context-dependent manner, with the RS domain contributing to binding specificity *in planta*, where At-RS31 undergoes post-translational modifications and interacts with other proteins. This highlights the importance of considering a combination of both *in vivo* and *in vitro* data to gain a more comprehensive understanding of the RNA-binding protein properties.

Alternative splicing is crucial in regulating At-RS31 and its paralogs, contributing to potential functional redundancy. This redundancy may explain why the overexpression line of *At-RS31* shows more extensive changes in gene expression than the mutant in our analysis. In natural context, this enables the paralogs to compensate for each other’s functions, maintaining stability in the regulatory network. Feedback regulation, where splicing and expression patterns of the paralogs are influenced by At-RS31 levels, indicates a tightly controlled balance in their activity. Similar cross-regulatory mechanisms were described for the paralogs of Arabidopsis glycine-rich proteins and polypyrimidine tract-binding proteins (Schöning *et al*., 2007;

Schoning *et al*., 2008; Stauffer *et al*., 2010; Burgardt *et al*., 2024). These mechanisms also occur in other taxa, where SR proteins and hnRNPs in mammals and Drosophila regulate each other’s expression through alternative splicing coupled with NMD, maintaining splicing homeostasis (Jumaa & Nielsen, 1997; Kumar & Lopez, 2005; Rossbach *et al*., 2009). This regulation, as well as the positive feedback loop between At-RS31 and At-SR30 and cross-regulation of SR proteins underscore the complexity of gene expression regulation in plants. At-RS31 shares 77% identity with the RRMs of its paralog At-RS31a, feature suggesting similar if not identical RNA binding preferences (Ray *et al*., 2013) and potentially the same target transcripts. Relatively high identity level, ∼62%, with the RRMs of At-RS40 and At-RS41 also suggest a potentially overlapping set of targets. Downregulation of the paralogs in the overexpression line and upregulation in the mutant suggests a robust mechanism of maintaining splicing homeostasis. On the other hand, changes in these RRMs across evolution potentially influenced their specificity towards different RNAs. Additionally, the paralogs have distinct RS regions suggesting differences in the array of interacting proteins as well as potential differences in the RNA recognition and their targets. Moreover, members of this subfamily show overlapping but distinct expression patterns in different plant organs, suggesting that they evolved to use distinct sets of targets (Kalyna 2004).

In previous research (Lopato *et al*., 1999), we demonstrated that At-SR30 overexpression modulates AS in *At-RS31*, leading to increase in its protein-coding isoform. The interplay between At-RS31 and At-SR30, where each enhances the production of the other’s protein-coding isoform, suggests a positive feedback loop. Furthermore, our recent studies indicate that protein-coding isoforms of both *At-RS31* and *At-SR30* increase in light and decrease in dark, a response mediated by chloroplast retrograde signaling (Petrillo *et al*., 2014), and show similar regulation in response to sugars and changes in TOR activity (Riegler *et al*., 2021). The potential feedback loop between At-RS31 and At-SR30, therefore, may be an important mechanism for amplifying the response to these changes, enabling the plant to optimize its functioning in fluctuating environmental conditions.

Interestingly, At-RS31 appears to be involved in regulation of the components of the ABA pathway and the core ABA-responsive genes (Finkelstein, 2013). ABA is a plant hormone regulating various aspects of plant growth and development and is essential for responses to a range of abiotic stresses. Increased ABA levels inhibit TOR signalling, thereby restricting growth under stress. Conversely, the TOR pathway counteracts ABA signalling to promote growth when stress is absent. This interplay provides a mechanism for balancing between plant growth and stress response (Wang *et al*., 2018). At-RS31 binding sites are found by iCLIP in 49 out of 313 genes. Moreover, these prevail in the core ABA-repressed genes (18 out of 47, ∼38%) (Table **S13**). At-RS31 modulates the expression of a wide range of ABA-related genes. Furthermore, At-RS31 binds and modulates the alternative splicing of the transcripts of key genes related to the TOR pathway. Given the regulation of At-RS31 by TOR (Riegler et al., 2021), At-RS31 may provide an additional regulatory layer that integrates ABA and TOR pathways. This integration could ensure that under stress conditions, ABA signalling is augmented to inhibit the TOR pathway, prioritizing stress responses over growth. Conversely, under favourable conditions, TOR signalling can regulate At-RS31 to support growth by modulating ABA signalling components. Consistently, the overexpression line of *At-RS31* has an ABA hypersensitive phenotype (Fig. **S13**).

In conclusion, by using iCLIP, RNAcompete and transcriptome analyses of *At-RS31* overexpression lines and mutants, we found that At-RS31 is a master splicing factor of key relevance to balance growth and stress responses in plants.

## ACKNOWLEDGEMENTS

The authors thank John Brown and Runxuan Zhang (The James Hutton Institute, Dundee, Scotland) for helpful discussions and providing information on the *A. thaliana* high-confidence transcription start sites. We thank Kristina Neudorf for expert technical assistance. We also acknowledge the Next Generation Sequencing Facility at Vienna BioCenter Core Facilities (VBCF), a member of the Vienna BioCenter (VBC), Austria, for performing RNA-seq, and the Genomics Core Facility at the Institute of Molecular Biology (Mainz, Germany) for sequencing the iCLIP libraries. This work was supported by grants from the Austrian Science Fund (FWF) (P26333 to MK), the German Research Foundation (DFG) (STA653/16-1 to DS), the Agencia Nacional de Promoción Científica y Tecnológica (ANPCyT, Argentina) (PICT 2020 02865 to EP), the International Centre for Genetic Engineering and Biotechnology (ICGEB) (CRP/ARG22-03 to EP), NIH R01 HG008613 to TH, CIHR Project Grant PJT-162255 and NIH R01 HG013328 to TH and QM, and NIH/NCI Cancer Center Support Grant P30 CA008748 to QM.

## AUTHOR CONTRIBUTIONS

MK, AB and DS conceived the study. MK, AB, DS, TH and QM acquired the funding. AF prepared the iCLIP and RNAcompete constructs and generated plants for iCLIP. TK performed the iCLIP experiments. ML performed the bioinformatics analyses of the iCLIP data. HZ expressed proteins for the RNAcompete assay and DR conducted the assay. YM prepared the RNA-seq libraries. PV and YM conducted the bioinformatics analyses of the RNA-seq data. BN, SF and SR carried out the RT-PCR experiments. MK, DS and EP critically analyzed the data and wrote the manuscript. All authors approved the final manuscript.

## DATA AVAILABILITY

RNA-seq reads have been deposited at the Short Read Archive under the accession numbers SRX2640702, SRX2640703, SRX2640704, SRX2640706, SRX2640707, SRX2640708, SRX2640710, SRX2640711, SRX2640712. iCLIP reads have been submitted to SRA under the BioProject accession PRJNA1179310.

## SUPPORTING INFORMATION

### SUPPLEMENTAL FIGURES

**Fig. S1**

At-RS31-GFP fusion protein expressed in transgenic plants used in the iCLIP

**Fig. S2**

GST-tagged At-RS31 fusion proteins used for RNAcompete

**Fig. S3**

Immunopurification of At-RS31 protein–RNA complexes from UV crosslinked *RS31::RS31-GFP* and *35S::GFP* plants and preparation of iCLIP libraries

**Fig. S4**

Genome-wide distribution of crosslink sites

**Fig. S5**

Sequence logo of At-RS31 binding sites enriched upstream of 5’ splice sites

**Fig. S6**

RNAcompete analysis of the At-RS31 protein

**Fig. S7**

RT-PCR analyses of differential alternative splicing events in genes with At-RS31 binding sites identified by iCLIP

**Fig. S8**

RT-PCR analyses of differential alternative splicing in genes encoding RNA binding proteins and splicing factors, including SR proteins

**Fig. S9**

Examples of At-RS31 and TOR pathway shared targets

### SUPPLEMENTARY TABLES

**Table S1**

Oligonucleotides used in this study

**Table S2**

iCLIP read statistics

**Table S3**

At-RS31 iCLIP binding site coordinates

**Table S4**

At-RS31 iCLIP target transcripts

**Table S5**

Functional enrichment analysis

**Table S6**

Distances from transcription start sites to At-RS31 binding sites

**Table S7**

Regions upstream of 5’ splice sites containing At-RS31 binding sites

**Table S8**

Differential alternative splicing analysis for At-RS31 mutant and overexpression plants

**Table S9**

Differential gene expression analysis for At-RS31 mutant and overexpression plants

**Table S10**

Transcription factors modulated by At-RS31

**Table S11**

Shared targets of At-RS31 and the TOR pathway

**Table S12**

At-RS31 in abscisic acid metabolism and signaling

