## Supporting Figures for "At-RS31 orchestrates hierarchical cross-regulation of splicing factors and integrates alternative splicing with TOR-ABA pathways"

(a)

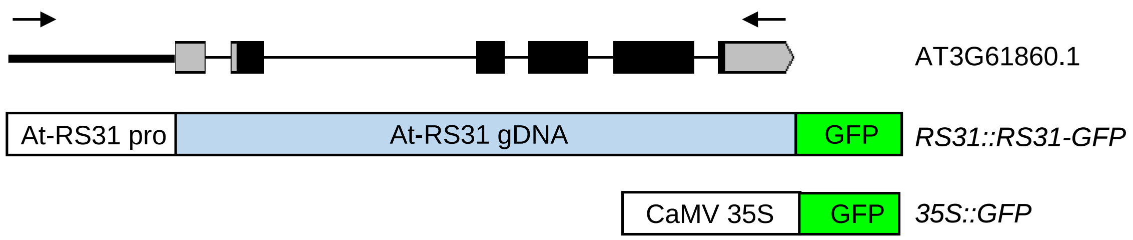


(b)

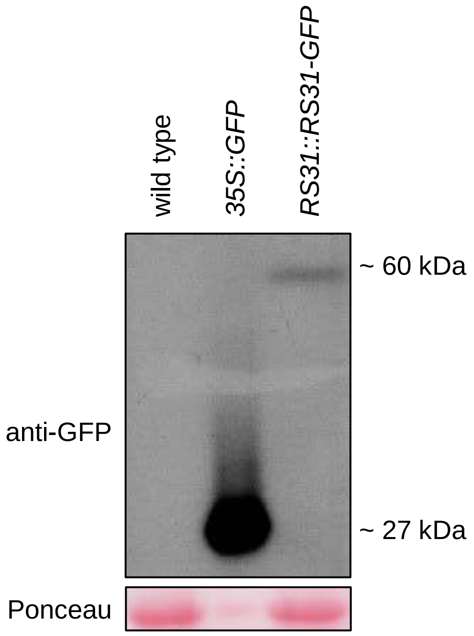


**Fig. S1**

At-RS31-GFP fusion protein expressed in transgenic plants used in the iCLIP

(a) *At-RS31* genomic region comprising the coding sequence as well as endogenous promoter, untranslated regions and introns was amplified by PCR (arrows – primers) and cloned upstream of the green fluorescent protein (GFP). AT361860.1 represents the reference gene model of *At-RS31*. Thick line – promoter region; grey and black boxes – exons in the untranslated and coding regions, respectively; thin lines – introns. GFP under the CaMV 35S promoter was used as a control. The constructs were used to generate *RS31::RS31-GFP* and *35S::GFP* transgenic plant lines used in the iCLIP.

(b) Immunoblotting of the wild type, *35S::GFP* and *RS31::RS31-GFP* plants using anti-GFP antibody. Wild type and *35S::GFP* plants were used as negative and positive controls, respectively. Ponceau S staining was performed to check loading. Note that only 1/10th of *35S::GFP* sample was loaded.

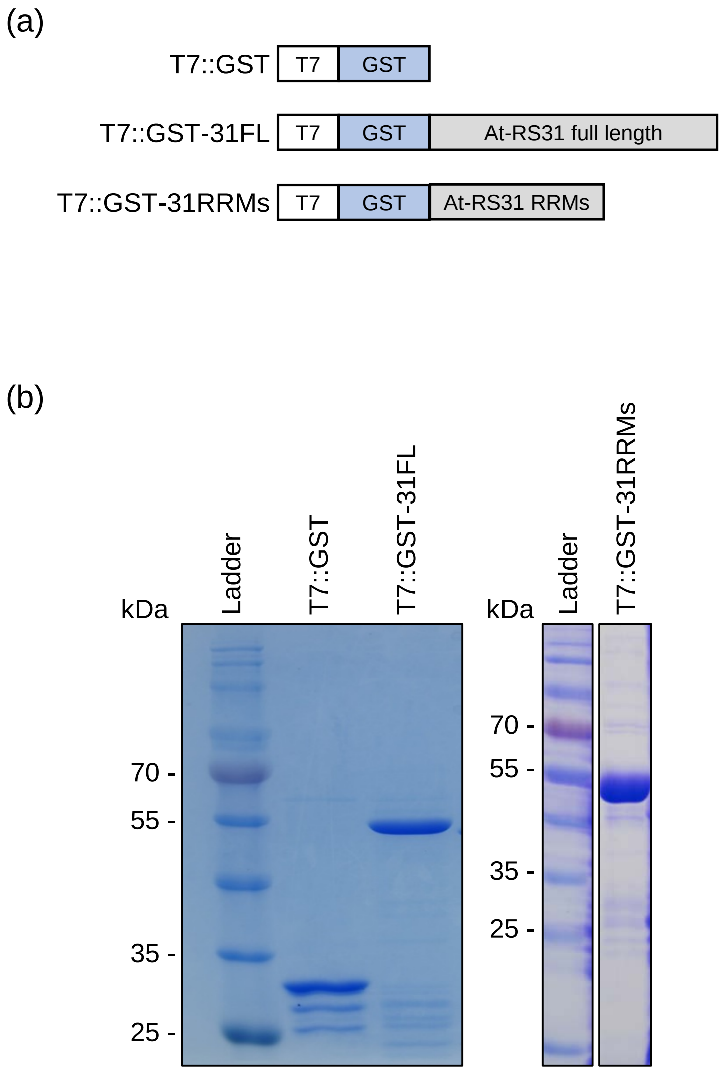


**Fig. S2**

GST-tagged At-RS31 fusion proteins used for RNAcompete

(a) N-terminal GST-tagged At-RS31 full length cDNA (T7::GST-31FL) and At-RS31 RRM (T7::GST-31RRMs) constructs.

(b) Purified GST and GST-tagged At-RS31 full length and RRMs-only fusion proteins were analysed by polyacrylamide gel electrophoresis and subsequent Coomassie staining. Marker positions are indicated.

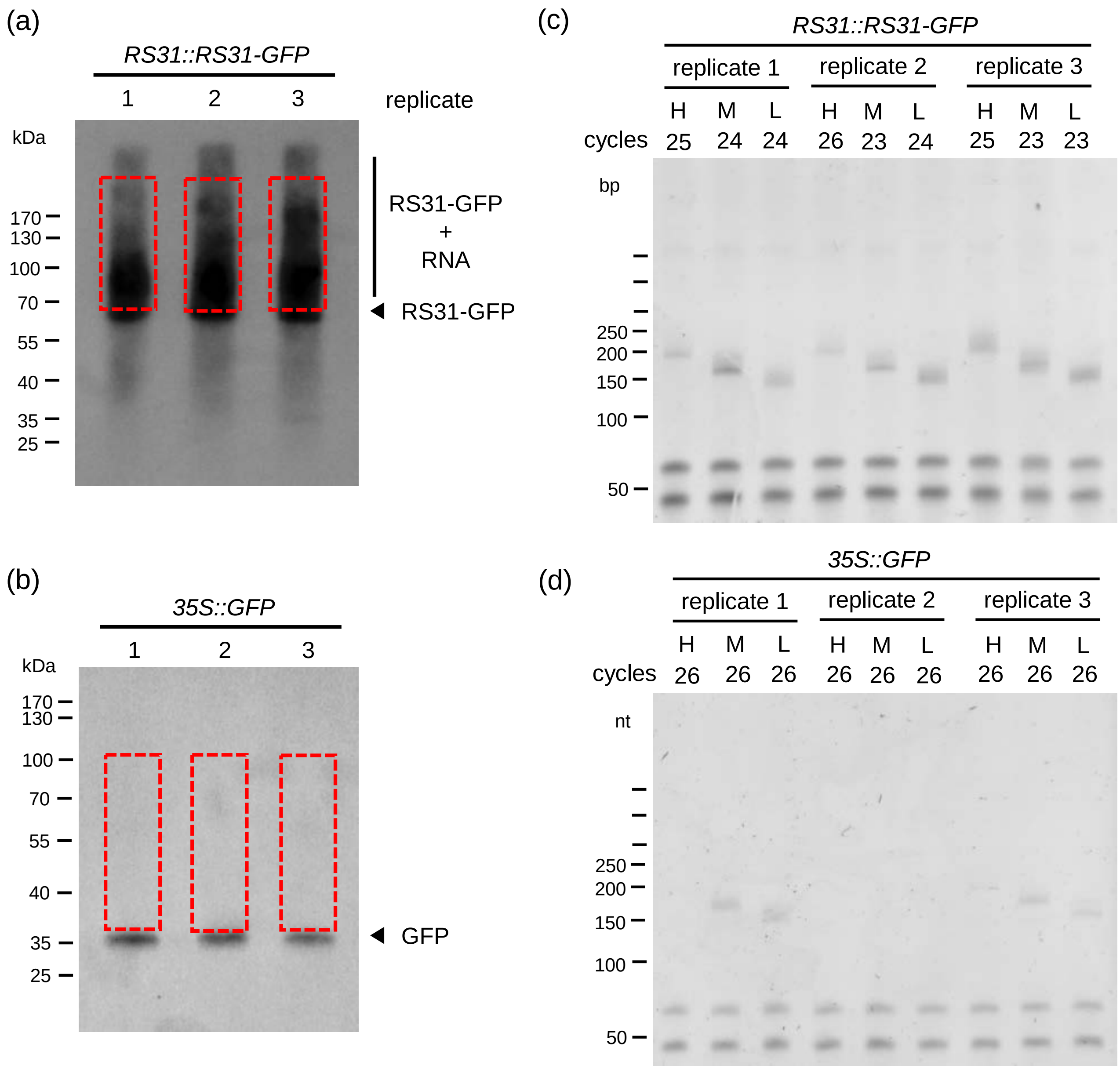


**Fig. S3**

Immunopurification of At-RS31 protein–RNA complexes from UV crosslinked *RS31::RS31-GFP* and *35S::GFP* plants and preparation of iCLIP libraries

(a, b) Autoradiograms of RS31-GFP (A) and GFP protein-RNA complexes (B) used for library preparation. The regions above the RS31-GFP and GFP proteins containing the co-precipitated RNAs, which were cut out for library preparation, are indicated by the red boxes. The positions of the markers are indicated.

(c, d) Preparation of iCLIP libraries by PCR amplification. The numbers of PCR cycles are indicated. The lengths of the cDNAs correspond to the size of the PCR product minus the length of the P3/P5Solexa primers and the barcode (128 nt in total). For each replicate, the low, medium and high molecular weight PCRs were combined according to their relative concentrations.

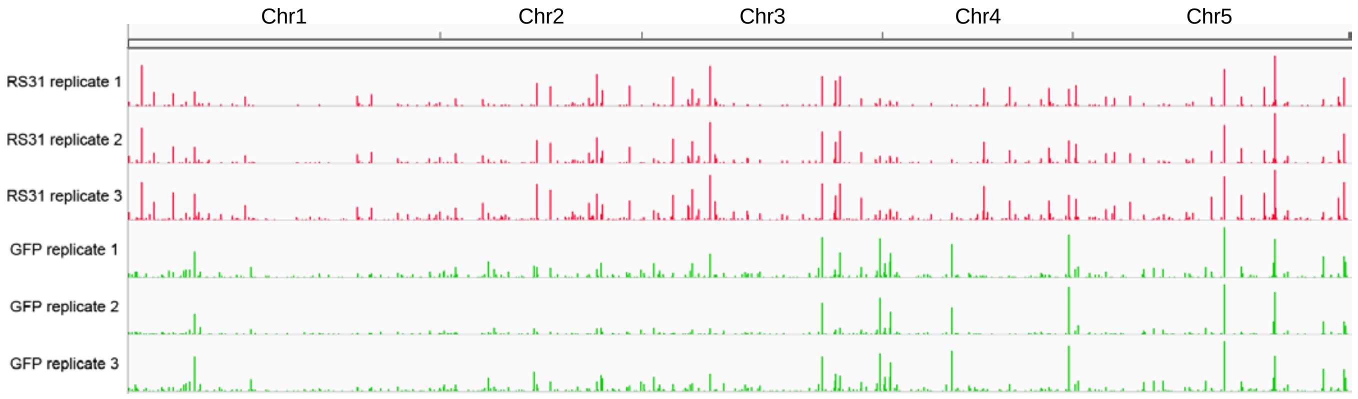


**Fig. S4**

Genome-wide distribution of crosslink sites

Localization of the crosslink sites of RS31-GFP (red) and GFP alone (green) throughout the *Arabidopsis thaliana* chromosomes for each biological replicate. The screenshot was created using the Integrative Genomics Viewer .

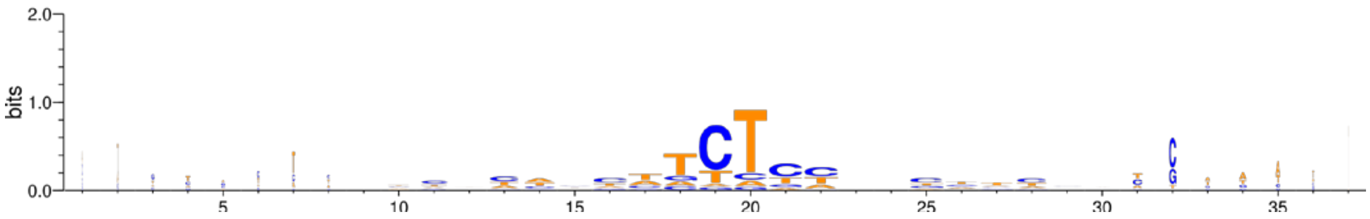


**Fig. S5**

Sequence logo of At-RS31 binding sites enriched upstream of 5' splice sites

Sequence logo from aligned RS31-GFP binding sites, which were enriched upstream of 5’ splice sites.

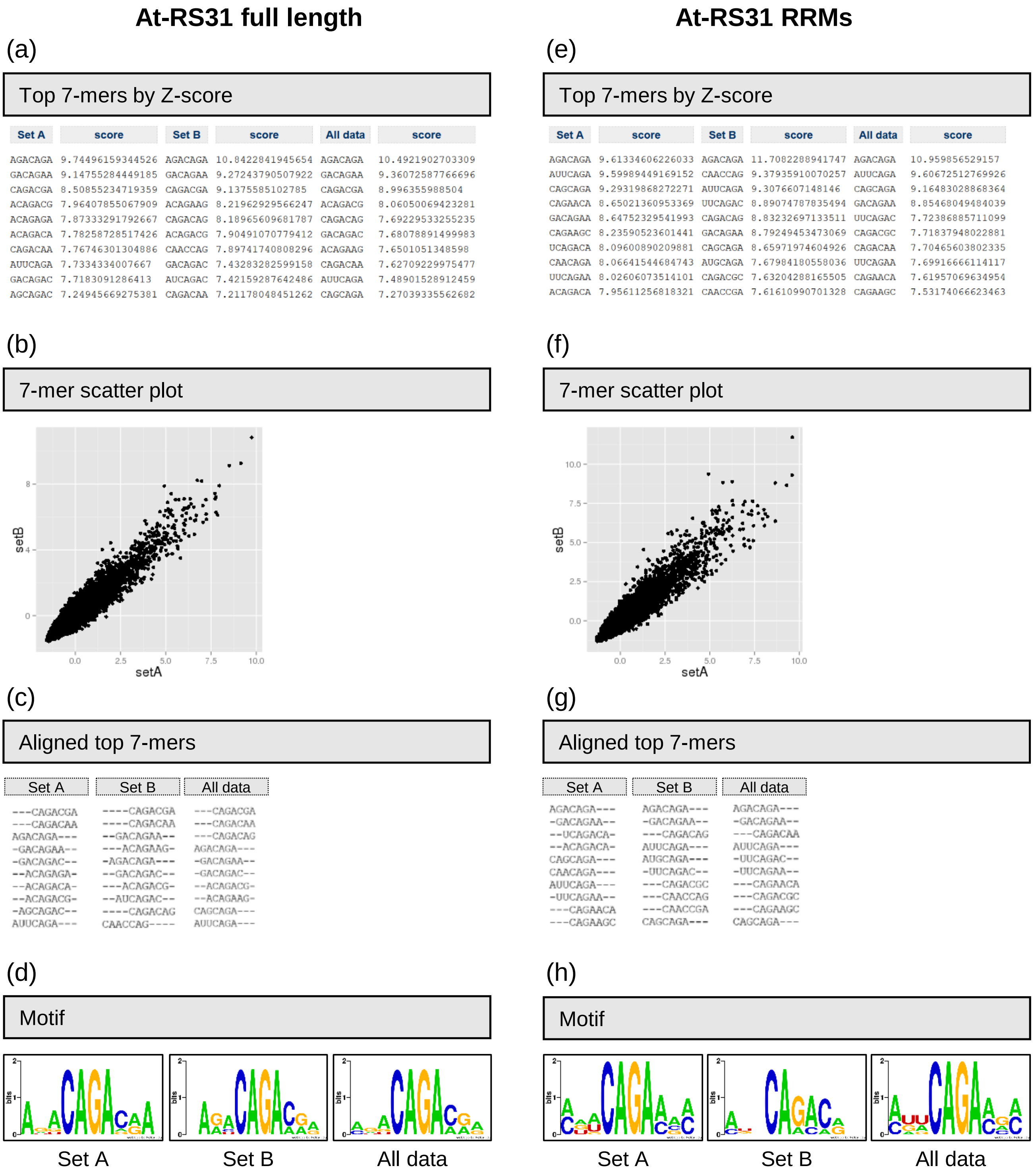


**Fig. S6**

RNAcompete analysis of the At-RS31 protein

RNAcompete assay was performed for the full-length At-RS31 protein (a-d) and its truncated version containing RRMs only (e-h).

(a) and (e) Top ten 7-mers for At-RS31 and At-RS31RRMs. Corresponding Z-scores for Set A, Set B, and the average of Set A and Set B (All data) are shown.

(b) and (f) Correlation between 7-mers Z-scores from Set A and Set B.

(c) and (g) Alignment of the top ten 7-mers for Set A, Set B, and Set A+B (All data).

(d) and (h) RNA-binding motifs of At-RS31 and At-RS31RRMs. Logos derived from the top ten 7-mers shown in (a) and (e). Sequence logos are generated by WebLogo at https://weblogo.berkeley.edu/.

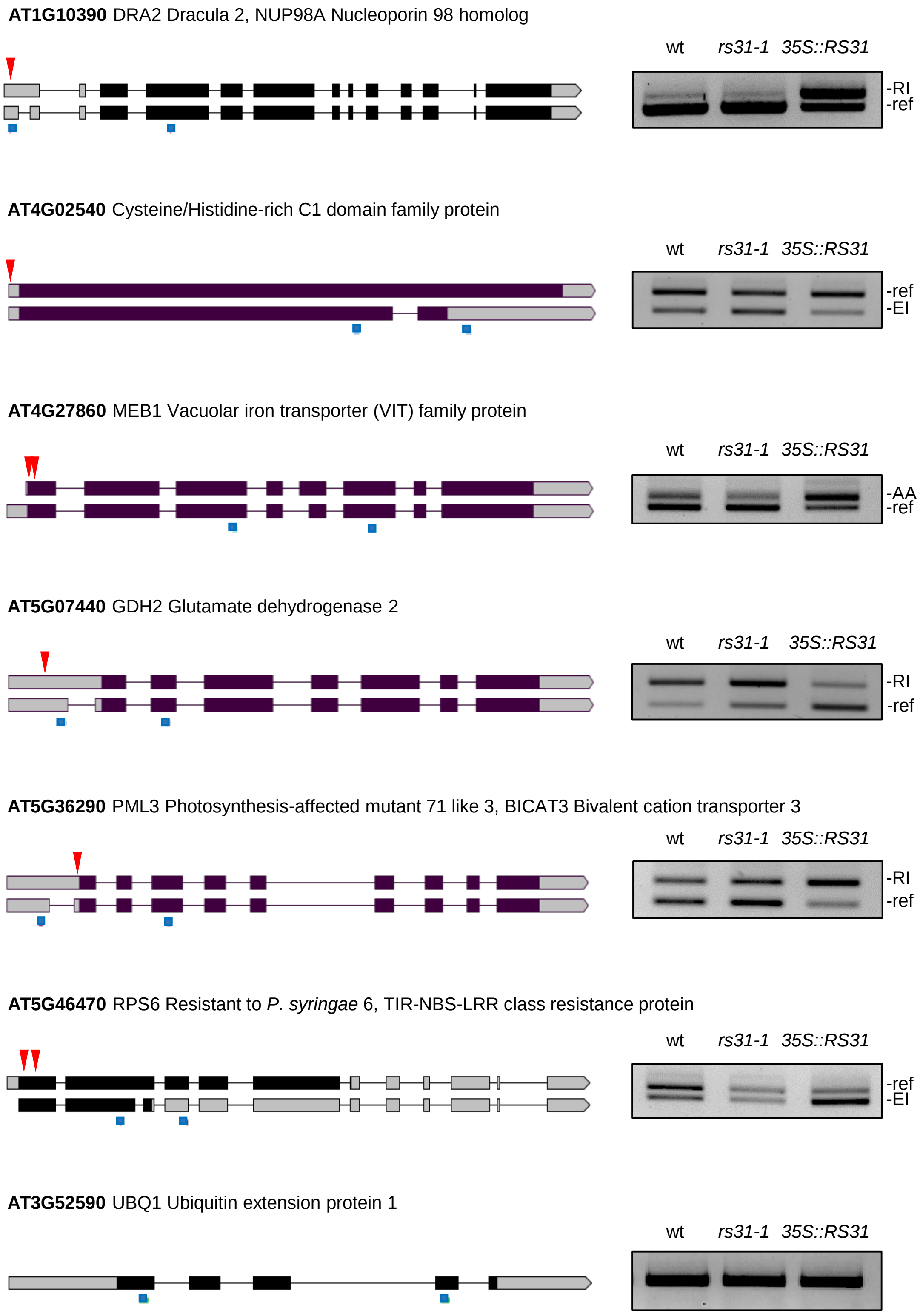


**Fig. S7**

RT-PCR analyses of differential alternative splicing events in genes with At-RS31 binding sites identified by iCLIP

RT-PCR analyses of differential alternative splicing (DAS) events in *rs31-1* mutant, *35S::RS31* overexpression and wild type (wt) plants were performed for genes with At-RS31 binding sites identified by iCLIP. Transcript models for the genes visualized using Boxify tool https://boxify.boku.ac.at/. Regions from translational start codon to stop codon or premature termination codon are shown in black. Red arrowheads show positions of At-RS31 binding sites. Blue squares below the transcript models represent primers used in the RT-PCRs, their positions were identified by Boxify. DAS events and reference (ref) transcripts encoding the full-length proteins are indicated to the right of the gels. RI - retained intron; EI – exitron; AA – alternative acceptor / alternative 3’ splice site. *UBQ1* was used as a loading control.

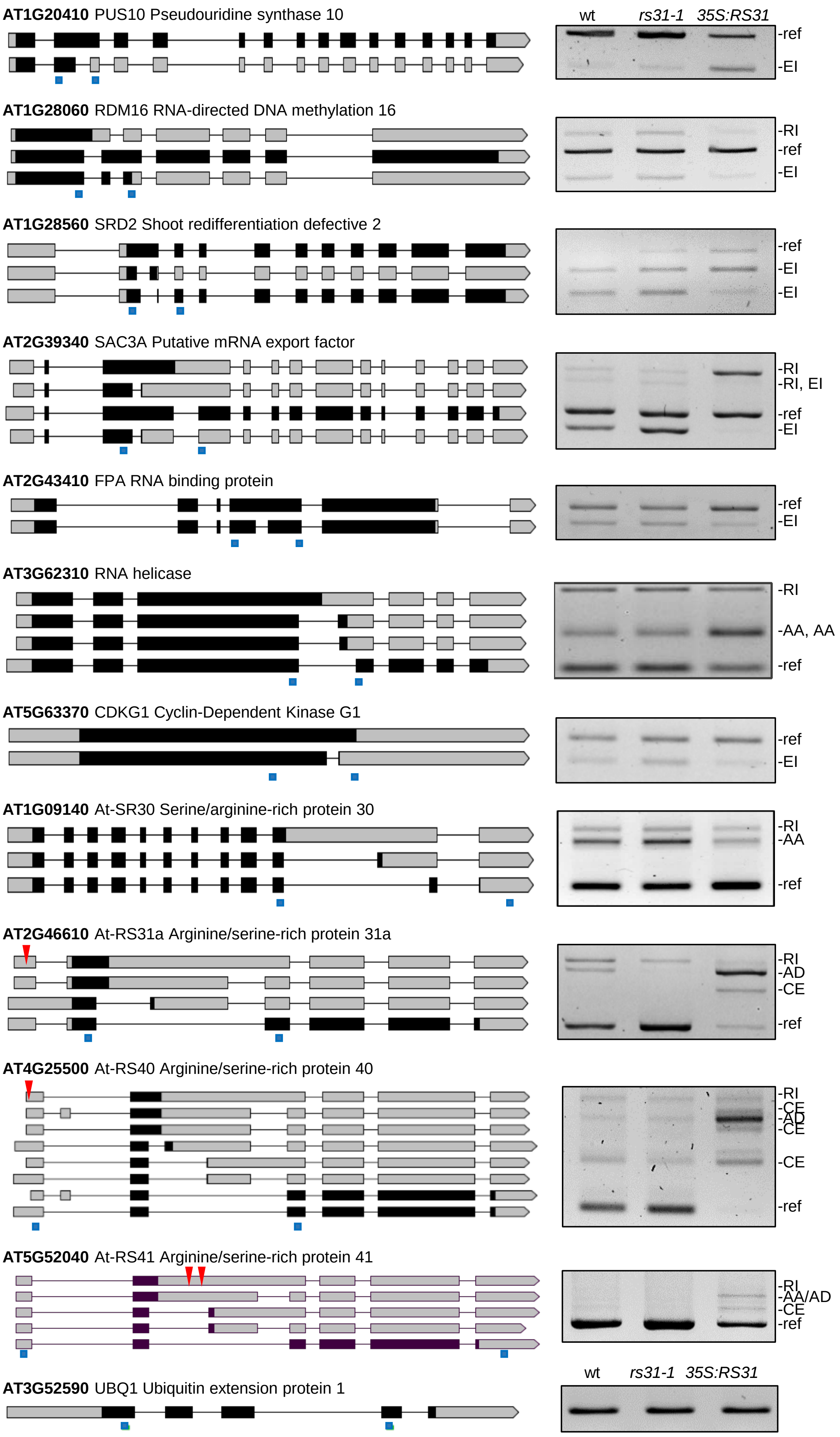
AT1G20410 PUS10 Pseudouridine synthase 10
wt rs31-1 35S:RS31
-ref
-EI
AT1G28060 RDM16 RNA-directed DNA methylation 16
-RI
-ref
-EI
AT1G28560 SRD2 Shoot redifferentiation defective 2
-ref
-EI
-EI
AT2G39340 SAC3A Putative mRNA export factor
-RI
-RI, EI
-ref
-EI
AT2G43410 FPA RNA binding protein
-ref
-EI
AT3G62310 RNA helicase
-RI
-AA, AA
-ref
AT5G63370 CDKG1 Cyclin-Dependent Kinase G1
-ref
-EI
AT1G09140 At-SR30 Serine/arginine-rich protein 30
-RI
-AA
-ref
AT2G46610 At-RS31a Arginine/serine-rich protein 31a
-RI
-AD
-CE
-ref
AT4G25500 At-RS40 Arginine/serine-rich protein 40
-RI
-CE
-AD
-CE
-CE
-ref
AT5G52040 At-RS41 Arginine/serine-rich protein 41
-RI
-AA/AD
-CE
-ref
wt rs31-1 35S:RS31
AT3G52590 UBQ1 Ubiquitin extension protein 1

**Fig. S8**

RT-PCR analyses of differential alternative splicing in genes encoding RNA binding proteins and splicing factors, including SR proteins

RT-PCR analysis was conducted to examine differential alternative splicing (DAS) events in *rs31-1* mutant, *35S::RS31* overexpression, and wild-type (wt) plants, focusing on genes encoding RNA binding proteins and splicing factors, including SR proteins. Transcript models for these genes are visualized using the Boxify tool (https://boxify.boku.ac.at/). Regions, spanning from the translational start codon to the stop codon or premature termination codon, are depicted in black. Blue squares beneath the transcript models represent the positions of primers used in the RT-PCR experiments, as determined by Boxify. Red arrowheads denote the locations of At-RS31 binding sites identified by iCLIP. DAS events and reference (ref) transcripts encoding the full-length proteins are indicated to the right of the gel images; AA – alternative acceptor / alternative 3' splice site; AD – alternative donor / alternative 5' splice site; CE – cassette exon / exon skipping; EI – exitron; RI - retained intron. The loading control used was *UBQ1*.

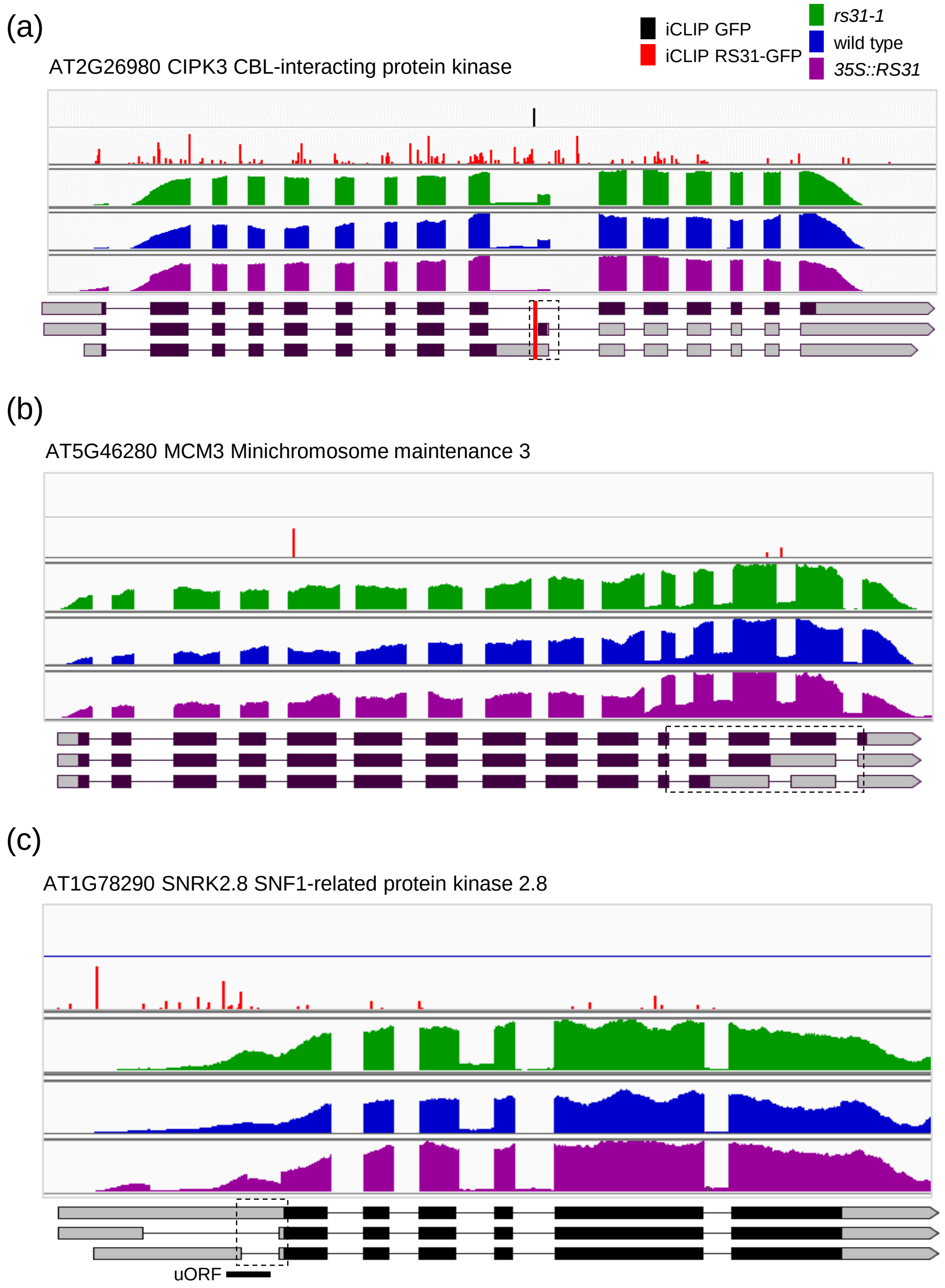


**Fig. S9**

Examples of At-RS31 and TOR pathway shared targets

(a) AT2G26980 CIPK3 CBL-interacting protein kinase; (b) AT5G46280 MCM3 Minichromosome maintenance 3; (c) AT1G78290 SNRK2.8 SNF1-related protein kinase 2.8

Integrated genome viewer (IGV) tracks are shown for RNA-seq read coverage in *rs31-1*, *35S::RS31* and wild type plants and for iCLIP crosslinks for RS31-GFP and GFP control. Transcript models are visualized using Boxify tool https://boxify.boku.ac.at/. Regions from translational start codon to the reference stop codon or a premature termination codon are shown in black. Dashed rectangles denote regions undergoing differential alternative splicing in *35S::RS31* or *rs31-1* in comparison to wild type plants. Vertical red lines show dominant binding peaks for At-RS31. uORF – upstream open reading frame.
