## Supporting Information for "At-RS31 orchestrates hierarchical cross-regulation of splicing factors and integrates alternative splicing with TOR-ABA pathways"

### SUPPLEMENTARY TABLES

**Table S1**

Oligonucleotides used in this study

**Table S2**

iCLIP read statistics

Table of read, called peak and binding site counts for each sample and processing step.

**Table S3**

At-RS31 iCLIP binding site coordinates

Table with genomic coordinates of RS31-GFP binding sites with the corresponding PureClip scores. Identifiers of associated protein-coding and non-coding (labeled NC) genes, as well as the position of the binding sites in the transcript regions (5´ UTR, CDS, intron, 3´ UTR) are indicated.

**Table S4**

At-RS31 iCLIP target transcripts

The gene annotation version was taken from Araport11, but only the reference gene models were considered.

**Table S5**

Functional enrichment analysis

Functional enrichment analysis of At-RS31 iCLIP targets, differentially alternatively spliced (DAS) and differentially expressed (DEG) genes in the *rs31-1* mutant and *35S::RS31* overexpressing plants has been performed using g:Profiler (Kolberg *et al.*, 2023) at https://biit.cs.ut.ee/gprofiler/gost. Version e110_eg57_p18_4b54a898. Term sources include the Gene Ontology (GO) for BP (biological process), MF (molecular function) and CC (cellular component) categories, as well as KEGG (Kyoto Encyclopedia of Genes and Genomes) pathways.

**Table S6**

Distances from transcription start sites to At-RS31 binding sites

**Table S7**

Regions upstream of 5' splice sites containing At-RS31 binding sites

Coordinates of RS31-GFP binding sites extended to 21 nucleotides, enriched upstream of 5' splice sites.

**Table S8**

Differential alternative splicing analysis for At-RS31 mutant and overexpression plants

Differential alternative splicing analysis was performed in the *rs31-1* mutant, *35S::RS31* overexpressing plants compared to wild-type (WT) controls using RNA-seq data. The table shows differential alternative splicing events obtained and quantified using Whippet. DAS events with a probability ≥ 0.9 and an absolute delta percent-spliced-in IPSII ≥ 0.1 were considered to be significant. AA - alternative acceptor, AD - alternative donor, CE – cassette exon / exon skipping, EI – exitron, RI - retained intron. The table includes gene expression differences in DAS genes based on the adjusted p-value < 0.05 (IL2FCI>1 not required) and classification of DAS genes as RNA-binding proteins or splicing factors (SF-RBP) or transcription factors (TF). At-RS31 iCLIP peak occurrences, their coordinates and location relative genic features are based on data shown in Tables S3 and S4.

**Table S9**

Differential gene expression analysis for At-RS31 mutant and overexpression plants

Differential gene expression analysis was performed in the *rs31-1* mutant, *35S::RS31* overexpressing plants compared to wild-type (WT) controls using RNA-seq data. A gene is significantly differentially expressed (DE) in a contrast group if it has adjusted p-value < 0.05 and |L2FC| ≥ 1. TPM - transcript per million reads. The table includes classification of DE genes as

RNA-binding proteins or splicing factors (SF-RBP) or transcription factors (TF). At-RS31 iCLIP peak occurrences in DE genes, their coordinates and location relative genic features are based on data shown in Tables S3 and S4.

**Table S10**

Transcription factors modulated by At-RS31

The table shows transcription factors (TF) undergoing differential alternative splicing (DAS) and differential expression (DE) in the *rs31-1* mutant, *35S::RS31* overexpressing plants compared to wild-type (WT) controls. TF gene list is based on Calixto et al. (2018). Details on DAS and DE in TF genes are shown in Tables S8 and S9. At-RS31 binding sites in DAS and DE TF genes are based on iCLIP data shown in Tables S3 and S4.

**Table S11**

Shared targets of At-RS31 and the TOR pathway

The table summarizes At-RS31 iCLIP binding sites and differential alternative splicing (DAS) events in the *rs31-1* mutant and *35S::RS31* overexpressing plants in genes related to the TOR pathway. It includes genes that show DAS under TOR inhibition (TOR DAS) (Riegler et al., 2021), genes encoding proteins regulated by TOR-dependent phosphorylation (TOR phosphoproteins) (Van Leene *et al.*, 2019; Scarpin *et al.*, 2020), and genes encoding proteins interacting with the TOR complex (TOR interactors) (Jamsheer K et al., 2022). ΔPSI - delta percent-spliced-in; AA – alternative acceptor / alternative 3' splice site; AD – alternative donor / alternative 5' splice site; CE – cassette exon / exon skipping; EI – exitron; RI - retained intron; CDS – coding sequence; UTR – untranslated region; NMD – nonsense-mediated mRNA decay; PTC – premature termination codon.

**Table S12**

At-RS31 in abscisic acid metabolism and signaling

The table summarizes At-RS31 iCLIP binding sites, differential alternative splicing (DAS) events, and differential expression (DE) in the *rs31-1* mutant and *35S::RS31* overexpressing plants in genes involved in metabolism or transport of ABA or signaling in response to ABA, as well as core ABA-induced or ABA-repressed genes (Finkelstein, 2013).
